# Dual spatio-temporal regulation of axon growth and microtubule dynamics by RhoA signaling pathways

**DOI:** 10.1101/2023.04.17.537156

**Authors:** José Wojnacki, Gonzalo Quassollo, Martín D. Bordenave, Nicolás Unsain, Gaby F. Martínez, Alan M. Szalai, Olivier Pertz, Gregg G. Gundersen, Francesca Bartolini, Fernando D. Stefani, Alfredo Cáceres, Mariano Bisbal

## Abstract

RhoA plays a crucial role in neuronal polarization, where its action restraining axon outgrowth has been thoroughly studied. We now report that RhoA has not only inhibitory but also a stimulatory effect on axon development depending on when and where exerts its action and the downstream effectors involved. In cultured hippocampal neurons, FRET imaging revealed that RhoA activity selectively localizes in growth cones of undifferentiated neurites, while in developing axons it displays a biphasic pattern, being low in nascent axons and high in elongating ones. RhoA-Rho kinase (ROCK) signaling prevents axon initiation but has no effect on elongation, while formin inhibition reduces axon extension without significantly altering initial outgrowth. Besides, RhoA-mDia promotes axon elongation by stimulating growth cone microtubule stability and assembly, as opposed to RhoA-ROCK that restrains growth cone microtubule assembly and protrusion. Finally, we show that similar mechanisms might operate during axonal regeneration, with RhoA-ROCK slowing axon regrowth after axotomy and RhoA-mDia favoring extension of regenerated axons.

## Introduction

Studies on cultured hippocampal and cortical neurons that develop *in situ* have established that the generation of an axon from a rather symmetric array of short and highly dynamic undifferentiated neurites is one of the early events underlying the establishment of morphological polarity (Caceres et al., 2012; Funahashi et al., 2014; Wilson et al., 2022). Current evidence favors the view that an increase in actin dynamics and growth cone size selects a minor neurite to become the axon (Bradke and Dotti, 1997, 1999; Kunda et al., 2001). It is believed that dynamic microtubules (MTs) penetrate the loose growth cone microfilament network allowing transport of polarizing and/or growth-promoting factors to the distal domain that in turn activate membrane-associated signaling pathways (Dent and Baas, 2014; Wojnacki et al., 2014) leading to MTs stabilization (Witte et al., 2008; Montenegro-Venegas et al., 2010). The newly formed axon would then initiate a phase of rapid elongation and branching. Not surprisingly, a considerable effort has been devoted to identify the molecular machinery involved in driving MT stability during axon formation (Conde and Caceres, 2009; van Beuningen and Hoogenraad, 2016). Nevertheless, key questions and mechanisms remain unsolved.

RhoA, a conspicuous small RhoGTPase family member (Hall and Lalli, 2010; Gonzalez-Billault et al., 2012), has been implicated in neuronal polarization. RhoA, and its downstream effector Rho Kinase (ROCK), exert an inhibitory tone preventing minor neurites from engaging in axon-like growth (Da Silva et al., 2003; Chuang et al., 2005; Takano et al., 2019); accordingly, suppression of Lfc, a RhoA guanosine-nucleotide exchange factor (GEF) promotes axon initiation while its up-regulation has the opposite effect (Conde et al., 2010; Wilson et al., 2020b). Inhibition of RhoA-ROCK signaling also impairs oriented axon formation *in situ* (Xu et al., 2015). Thus, inactivation of RhoA appears to be sufficient to trigger axon formation and hence the establishment of neuronal polarity. However, a recent study has provided evidence suggesting that while RhoA restrains axon initiation and growth, it has no effect on axon specification (Dupraz et al., 2019).

Another set of evidence suggests a more complex function of RhoA during axonogenesis. For example, in cultured cerebellar granule neurons, high RhoA activity downstream of stromal cell-derived factor-1α (SDF-1α) inhibits axonal elongation through ROCK. Surprisingly mild or low RhoA activity was reported to have the opposite effect promoting axonal elongation through a different effector, namely mDia1 (Arakawa et al., 2003). A dual effect has also been reported in NGF-stimulated PC12 cells, where, RhoA activation inhibits neurite outgrowth but, if activated afterwards, it promotes neurite elongation (Sebok et al., 1999). These results suggest that the function of RhoA is regulated both spatially and temporally. So far, the spatio-temporal activation pattern of RhoA during neuronal polarization remains unknown. There is no information regarding distinct RhoA effectors regulating different stages of axon development or over different neuronal compartments. Additionally, RhoA regulates MT stability/dynamics in other systems, but has not been tested for regulating MT dynamics/stability during axon growth,

Here, we report evidence of dual role of RhoA signaling during de novo and regenerative axonal growth. First, using FRET-based biosensors (Fritz et al., 2013; Li et al., 2017) we found that RhoA is highly active in growth cones of minor neurites and elongating axons; by contrast, low growth cone RhoA activity parallels the transformation of a minor neurite into an axon. Interestingly, ROCK activity is high in minor neurites, but low in both sprouting and elongating axons. Lysophosphatidic acid, a strong RhoA activator, prevents axon formation in stage 2 neurons (e.g. neurons with several minor neurites but no axon), but enhances elongation in neurons with established axon (e.g. stage 3 neurons); accordingly, RhoA suppression enhances axon outgrowth in early polarizing neurons but reduces axonal elongation in stage 3 polarized neurons. We found that diverging signaling downstream of RhoA is achieved by means of different effector proteins acting on MTs dynamics. While ROCK mediates RhoA’s inhibitory effects, mDia stimulates axonal extension by stabilizing MTs in the axonal growth cone. Finally, we show that similar mechanisms might operate after axotomy, with RhoA-ROCK slowing axon regrowth and RhoA-mDia, favoring regenerative axon elongation.

## Results

### RhoA activity in minor processes and axons during neuronal polarization

In the first set of experiments, we used a 1^st^ generation RhoA-FRET biosensor (RhoA1G), based on cyan fluorescent protein (CFP) as donor and yellow fluorescent protein (YFP) as acceptor (Supplementary Fig. 1, see also Fritz et al., 2013), to examine RhoA activity during the so-called second (growth) phase of polarization (Caceres et al., 2012), when a single axon is generated from an array of multiple short and highly dynamic minor neurites.

To this end, neuronal cultures were transfected 2, 18, 24 and 48 h after plating with RhoA1G, fixed 18 h later and RhoA activity evaluated by FRET using ratiometric imaging. We validated the FRET assay using CHO cells transfected with constitutive active (Q63L), dominant negative (T19N) and wild type (WT) versions of the RhoA biosensor (Supplementary Fig. 1 *A-C*). In agreement to previous reports with this biosensor (Fritz et al., 2013), our measurements yielded an average relative-FRET difference between Q63L and T19N of ∼20% (Supplementary Fig. 1D). In addition, the mutant biosensors showed no discernible spatial activation patterns while the WT variant displayed the highest RhoA FRET signal at the leading edge of the cells (Supplementary Fig. 1C, arrows), similar to what has been described for randomly migrating <u>m</u>ouse <u>e</u>mbryonic fibroblasts (MEFs; (Machacek et al., 2009)).

A similar analysis performed in 1-3 DIV (days in vitro) hippocampal neurons, revealed that high RhoA activity was spatially confined to specific cellular domains (Fig. 1), despite that RhoA1G YFP-fluorescence was found homogenously distributed throughout the neuron, including the cell body, the neurites, and their growth cones (Supplementary Fig. 2, see also (Gonzalez-Billault et al., 2012)). A representative image showing color-coded RhoA activity before axon formation (multipolar stage-2 neurons) is shown in Fig. 1*A*. In such neurons (∼20 h i*n vitro*) high RhoA activity was predominantly detected at tips (Fig. 1*A*, arrowheads) or growth cones (Fig. 1*A*, arrows) of minor processes. Remarkably, this pattern of RhoA activity changes dramatically upon onset of the transition from stage 2 to stage 3; i.e. during axon sprouting. The transformation of a minor neurite into an axon has been associated with an increase in growth cone size, accumulation of growth promoting factors and reorganization of the MTs and actin cytoskeletons (Bradke and Dotti, 1997, 1999; Kunda et al., 2001; Witte et al., 2008; Caceres et al., 2012). A considerable decrease of RhoA activity was detected in the enlarged growth cones (Fig. 1*B*, Supplementary Fig. 2A, B), concomitant with high Rac1 (Fig. 1*C*, arrow; Supplementary Fig. 2C, D) and Cdc42 (Chuang et al., 2005) activities.

**Figure 1.**
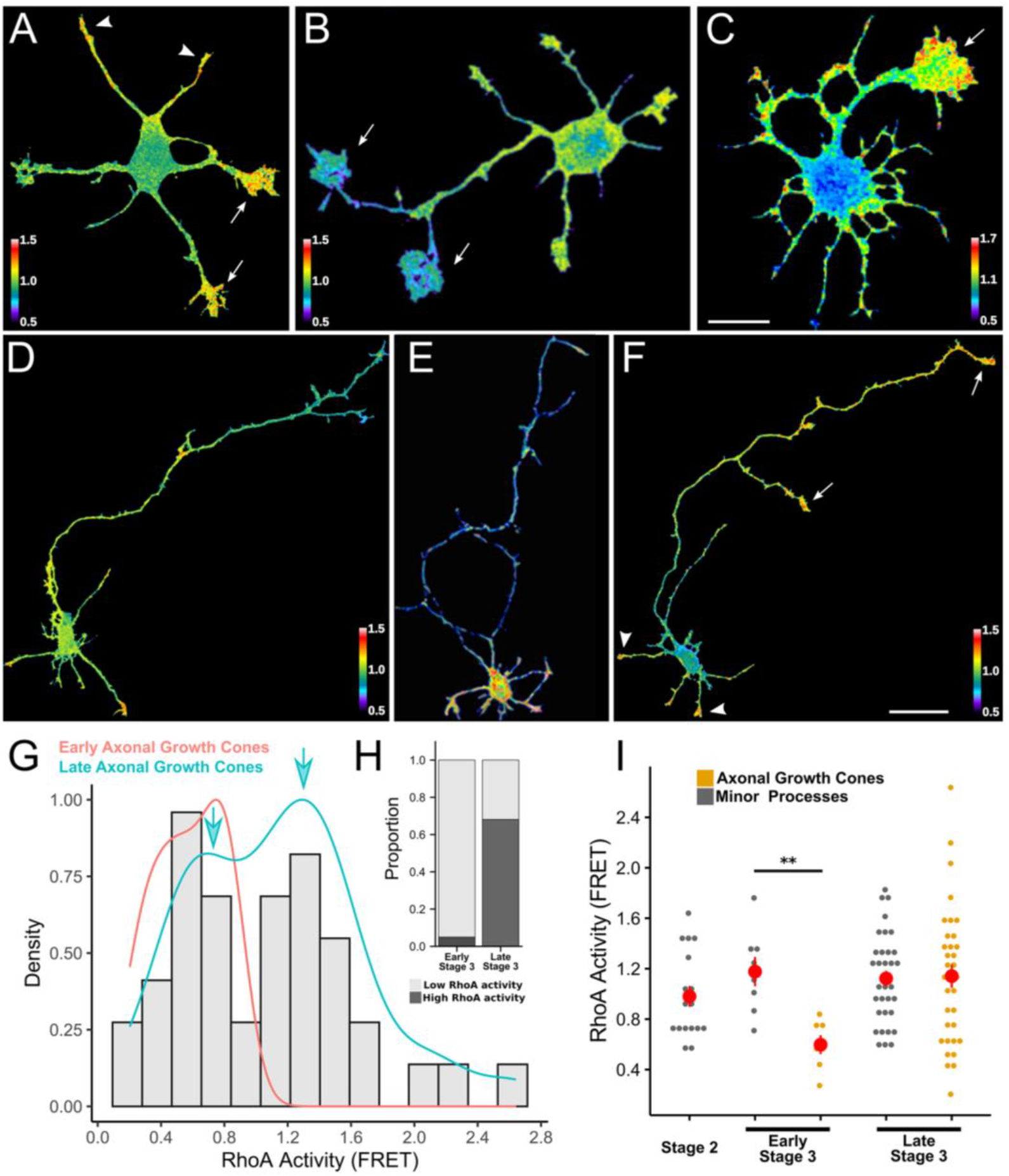
RhoA activity patterns during neuronal differentiation. (*A*-*C*) Representative FRET map images showing RhoA and Rac1 activities in 20 h cultured hippocampal neurons. FRET measurements were performed using the unimolecular RhoA1G (*A* and *B*) and Raichu-Rac1 biosensors (*C*). Note the difference in RhoA and Rac1 activities in the large growth cones of future axons (white arrows). Scale bar: 10 µm. (*D*-*F*) Representative FRET map images showing three different examples of RhoA activity in 36 h cultured hippocampal neurons. FRET measurements were performed using the RhoA1G biosensor. Note that there are different patterns of spatial RhoA activity including a subset of neurons with high RhoA activity in the distal axonal shaft and its growth cone (white arrows in *F*) beside in minor process growth cones (white arrowhead in *F*). Scale bar: 20 µm (*G*) Graph showing the frequency histogram of RhoA activity in stage 3 axonal growth cones. Bars are the density histogram of RhoA activity (FRET) in early and late axonal growth cones (n= 82). The choice of the density histogram allows us to compare the distribution of RhoA activity in early (pink line, n=14) and late (cyan line, n=68) axonal growth cones, which have different numbers of observations. Both lines are smoothed versions of the histogram separated as early or late stage 3. Both density lines were scaled to the same height. Note that early axonal growth cone (pink density line) falls almost exclusively in the low RhoA activity area while late ones fall in 2 distinct activity groups, low and high (cyan arrows). (*H*) Graph showing the proportion of early and late stage 3 neurons, displaying low and high RhoA activity in axonal growth cones; these values were calculated from 4 independent cultures. (*I*) Graph showing quantification of RhoA activity in minor process (grey) and axonal (orange) growth cones of early and late stage 3 neurons. Each small dot represents the FRET value in the growth cone of a single neuron. Each large red dot represents the mean ± S.E.M.; **<0.01; one-way ANOVA and Tukey’s *post hoc* test. FRET maps are color-coded according to activation intensity.

Low RhoA activity was also detected in more than 80% of newly formed axons (early stage 3; 24-30 h *in vitro*), however, by 36 h *in vitro* it was possible to distinguish axonal populations similar in length but with differences in RhoA activity (Fig. 1*D*-*G*). Interestingly, at later time points (late stage 3), when most of the axons become longer than 150 µm and begin to extend collateral branches, the percentage of cells displaying low axonal RhoA activity decreased significantly (Fig. 1*H*), with almost 70% of them showing a high RhoA FRET signal along the distal axonal segment including the growth cone. Quantitative analysis revealed that axonal growth cones of early stage 3 neurons have on average a 50% decrease in RhoA activity compared to equivalent ones of minor processes of the same cell or of stage 2 neurons (Fig. 1*I*). It also revealed that RhoA activity in axonal growth cones of late stage 3 neurons increases significantly becoming similar to that of the remaining minor neurites of the same neuron or of stage 2 neurons (Fig. 1*I*). By 3 DIV, almost all axons (∼300 µm in length) displayed high RhoA activity at their tips and/or along the distal third of the axonal shaft.

### RhoA has both inhibitory and stimulatory effects on process extension

The FRET experiments showed that active RhoA could be found in neurites that exhibit little if any net growth, such as minor processes of either stage 2 or stage 3 neurons, and in elongating axons of stage 3 neurons. These observations raised the possibility that in actively growing axons RhoA might act as an inhibitory brake to prevent excessive growth or alternatively that it has a stimulatory effect.

To begin testing this last idea we first evaluated the effect of the acute down-regulation of RhoA on axon growth by the use of an RNAi targeting RhoA ((Dupraz et al., 2019); see Methods). To examine RhoA function on initial axon outgrowth, freshly plated neurons were electroporated with the RNAi-*RhoA* at the time of plating and analyzed 1 day later. In agreement with Dupraz et al. (Dupraz et al., 2019), the early suppression of RhoA led to neurons displaying longer Tau-1 + axon-like neurites when compared with control cells (Fig. 2A-*C*). Also, it produced an increased percentage of neurons reaching stage 3 of neurite development by 1 DIV (Fig. 2D). A similar phenomenon was observed when neurons were treated 4 h after plating and analyzed 1 day later with C3-toxin, a well-known RhoA inhibitor ((Tsuge et al., 2015); Supplementary Fig. 3 *A*-*D*). By contrast, when the RNAi-*RhoA* treatment was applied 2 days after plating and neurons examined 24 or 48 h later, a significant decrease in axonal length was detected (Fig. 2E-*G*). Interestingly, such effect was not observed in minor processes, which are destined to become dendrites (Fig. 2H).

**Figure 2.**
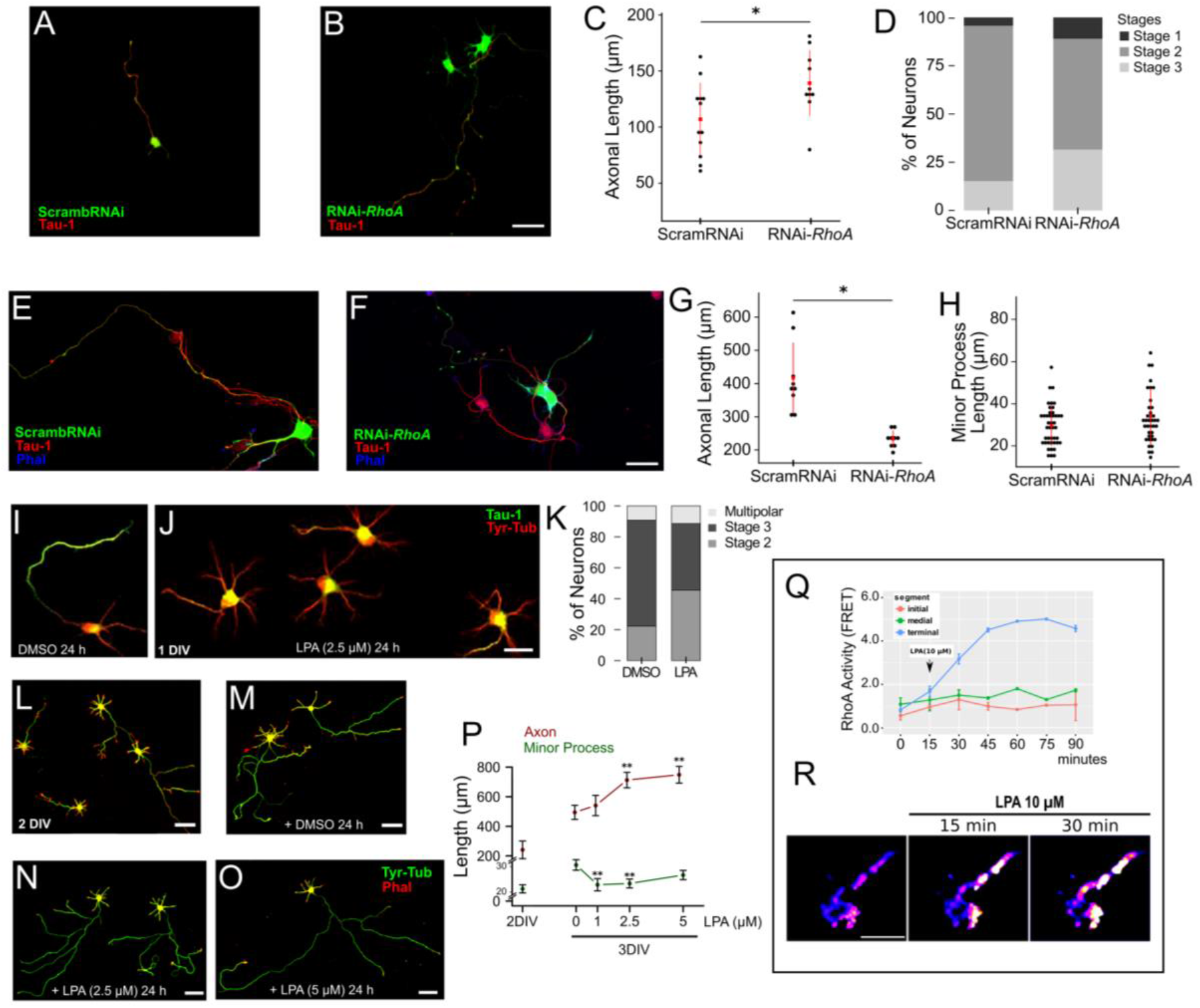
RhoA have opposite effects in axonal growth during neuronal polarization. (*A*, *B*) Representatives confocal images of cultured hippocampal neurons (1 DIV) co-electroporated with scrambleRNAi + GFP (*A*) or RNAi-*RhoA* + GFP (*B*) at the time of plating, fixed 24 h later and stained with the anti-Tau-1 mAb (red). Scale bar: 20 µm. (*C*) Graph showing quantification of axonal length in control (scramble-RNAi) or RNAi-*RhoA*-treated neurons. Each black dot represents the axonal length of a single neuron. Red dots represent the mean total axonal length ± S.E.M.; *<0.05; Student’s t-test. For all experiments, 12 neurons were quantified and pooled from at least three independent cultures. (*D*) Graph showing percentage of stages 1, 2 and 3 in scrambleRNAi or RNAi-*RhoA* expressing neurons. The percentages were calculated from 3 independent cultures. (*E*, *F*) Confocal images showing representatives examples of cultured hippocampal neurons (3 DIV) transfected with scrambleRNAi + GFP (*E*) or RNAi-*RhoA* + GFP (*F*). Cultures were transfected at 2 DIV, fixed 24 h later and stained with the anti-Tau-1 mAb (red) and Phalloidin-Alexa 633 (blue). Scale bar: 20 µm. (*G*, *H*) Graphs showing quantification of total length of Tau-1 positive axons (*G*) and minor process (*H*). Each black dot represents the neurite length of a single neuron. Red dot represents the mean neurite length ± S.E.M.; *<0.05; one-way Student’s t-test. For all experiments, 15 neurons were quantified pooled from at least three independent cultures. (*I*, *J*) Representatives confocal images of cultured hippocampal neurons treated with vehicle (DMSO, *I*) or with LPA (2.5 µM, *J*) for 24 h, fixed and immunostained for Tau-1 (green) and tyrosinated α-tubulin (red) Scale bar: 20 µm. (*K*) Graph showing the percentage of stages 2, 3 and multipolar in vehicle (DMSO) or LPA treated neurons. The percentages were calculated from 3 independent cultures. (*L*-*O*) Representative confocal images of cultured hippocampal neurons before (2 DIV, *L*) and after 24 h of LPA treatment (vehicle, *M*; 2,5 µM, *N* and 5 µM, *O*). Neurons were stained with mAb Tyrosinated α-tubulin (green) and phalloidin-rhodamine (red). Scale bar: 20 µm. (*P*) Graphs showing quantifications of average total length of minor (green) and axonal (red) processes of 3DIV hippocampal neurons after 24 h treatments with different doses of LPA. Graphs represent mean ± S.E.M.; **<0.01; one-way ANOVA and Tukey’s *post hoc* test. For all experiments and each condition, 12 to 25 neurons were quantified and pooled from at least three independent cultures. (*Q*) Time course analysis of ratiometric FRET RhoA activity at initial, medial and terminal segments of 3 DIV axons before and after treatment with LPA (10 µm, arrow). Note the marked increase of RhoA activity in terminal axonal segment after the LPA treatment. Graphs represent mean ± S.E.M. (*R*) Time lapse images showing the effect of LPA (5 µM) on RhoA activity at the distal end of an axon from a stage 3 hippocampal neuron; note the considerable increase in RhoA activity after 15 min. Scale bar: 5 µm.

To further validate these observations, we also analyzed the effect of lysophosphatidic acid (LPA), a well-known RhoA activator (Li and Gundersen, 2008; Gonzalez-Billault et al., 2012; Choi and Chun, 2013; Quassollo et al., 2015), on neurite formation/extension at different stages of neuronal development. In the first group of experiments LPA was added to the culture medium 4-6 h after plating. After 1 day of treatment the cultures were fixed and stained for Tau-1 and tyrosinated α-tubulin (Tyr-Tub) (Fig. 2I-*K*). LPA (2.5 µM) treatment resulted in a significant decrease (almost 40%) in the number of neurons capable of sprouting an axon (Fig. 2K). Halted cells failed to localize Tau-1 to the distal third of a single neurite/axon, a typical feature of polarized neurons (Fig. 2I, J); instead, most Tau-1 immunofluorescence was found at the cell body (Fig. 2J). In addition, all these cells lack large growth cones distinctive of prospective axons. A 10 µM LPA dose had a more profound effect with more than 90% of the cells failing to develop an axon and with minor neurites significantly shorter than equivalent ones from control neurons.

To further test our hypothesis we evaluated the effect of LPA in older cultures (2 DIV), when more than 80% of cells have reached stage 3 of neurite development. When such neurons were treated for 24 h with different doses of LPA, axonal length increased significantly compared with control neurons (Fig. 2L-*O*). The highest concentration tested (5.0 µM) increased axonal length as much as 40% above the value of the control group (Fig. 2P). By contrast, all tested LPA doses (1.0, 2.5 and 5.0 µM) significantly decreased the length of minor neurites compared with non-treated ones (Fig. 2P). A similar response was observed when the LPA treatment was initiated at 3 DIV and analyzed 1 day later (not shown). Finally, we also observed that in 2 or 3 DIV neurons C3 toxin significantly reduced axonal length and LPA-stimulated axonal growth (Supplementary Fig. 3 *E*-*I*). FRET experiments revealed that LPA treatment (5.0 or 10.0 µM) significantly increased RhoA activity in the distal part of the axon, including their growth cones (Fig. 2Q, R). Typically, this increase occurred within 30 min and lasted for at least 90 min, the last time point analyzed (Fig. 2Q, R).

Together, these observations support the existence of a RhoA inhibitory tone on axon outgrowth (Caceres et al., 2012; Dupraz et al., 2019; Takano et al., 2019) but more importantly reveal for the first time that once axons are formed RhoA activity shifts this negative influence to become a promoter of axon elongation. The latter results raise the possibility that distinct effector pathways operate at different developmental stages (e.g. axon outgrowth vs. axon elongation) and/or cell domains (minor neurites vs. axons).

### ROCK regulates axon outgrowth but not elongation

Rho-kinase (ROCK) is a RhoA effector with a wide variety of functions including regulation of motility, morphology and neuronal polarity (Amano et al., 2010; Caceres et al., 2012; Takano et al., 2017; Takano et al., 2019). As expected, inhibition of ROCK with Y27632 enhanced axonal growth when applied 6 h after plating and examined at 1 DIV (Fig. 3A-*B*). This treatment also increased the percentage of neurons with several axon-like (Tau-1 +) neurites or multipolar neurons (Fig. 3C-*D*) and reverted the inhibitory effect of LPA on axon outgrowth (Fig. 3D).

**Figure 3.**
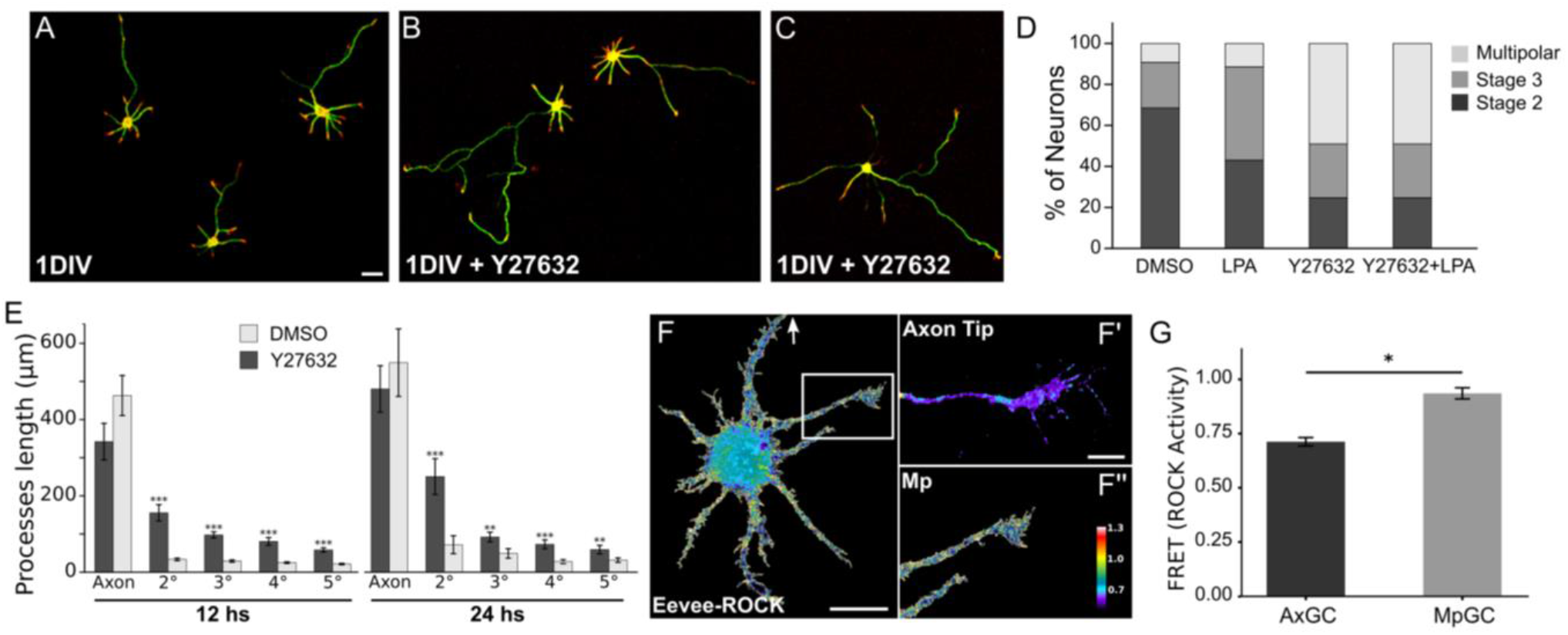
The RhoA effector ROCK promotes axon outgrowth but not elongation. (*A*-*C*) Representative confocal images of cultured hippocampal neurons (1 DIV) treated with vehicle (DMSO, *A*) or with Y27632 (10 µM, *B*-*C*) at the time of plating, fixed 24 h later and stained with an anti-tyrosinated α-tubulin mAb (green) and rhodamine-phalloidin (red). Scale bar: 20 µm. (*D*) Graph showing the percentage of stage 2, stage 3 and multipolar neurons in cultures treated for 24 h with vehicle (DMSO), LPA (10 µM), Y27632 (10 µM) or Y27632 + LPA (10 µM each). Percentages were calculated from 3 independent cultures. (*E*) Graphs showing quantification of average total axonal or minor processes (2°-5°) length of 3DIV neurons treated for 12 (left) or 24 (right) h with vehicle (DMSO) or Y27632 (10 µM). Note that the ROCK inhibitor increases minor processes length without affecting that of the axon. Neural processes were classified according to length. Graphs represent mean ± S.E.M.; **<0.01, ***<0.001; Student’s t-test. For all experiment, 12 to 20 neurons were quantified and pooled from at least three independent cultures. (*F*) Representative FRET map image showing ROCK activity in a hippocampal neuron from a 3DIV culture. FRET measurements were performed using the unimolecular Eevee-ROCK FRET-based biosensors. Scale bar: 20 µm. Ratiometric method was used to measure the FRET signal and FRET maps are color-coded according to the activation intensity. Color-code according to the scale bar. (*F*’-*F*’’) High magnification views of the axonal tip (arrow in *F*) and a minor process (Mp, insert in *F*) showing ROCK activity (*F*). Scale bar: 5 µm. Note that ROCK activity is very low at the distal end of the axon including its growth cone. (*G*) Graph showing quantification of ROCK activity in axonal (AxGC) and minor process (MpGC) growth cones of 3 DIV neurons. Graphs represent mean ± S.E.M.; *<0.05; Student’s t-test. For all experiment, 16 neurons were quantified and pooled from at least three independent cultures.

To determine if ROCK continues operating at later stages of axon development (e.g. during axon elongation) 2 DIV cultures were treated with Y27632 for different time periods. Neurites were arranged according to their length from longest to shortest, always defining the axon as the longest one (total length of at least 150 µm) (Fig. 3E). In these neurons, a 12 or 24 h treatment with Y27632 caused a 2-to 3-fold increase in minor neurite length compared with control cultures (Fig. 3E). Remarkably, the length of the differentiated axon (the longest neurite) was not significantly increased by Y27632 treatment (Fig. 3E). Thus, ROCK is neither inhibiting nor promoting axon elongation once the axon has been formed and is elongating. One possible explanation could be that elongating axons have low ROCK activity. To test this idea, we used Eevee-ROCK, a FRET-based biosensor with high sensitivity and specificity for ROCK activity (Li et al., 2017); see also (Takano et al., 2017). FRET revealed that ROCK activity is restricted to shafts and growth cones of minor neurites (Fig. 3F-*G*), declining progressively towards the distal axonal end. The lowest ROCK activity was detected in the axon growth cone (Fig. 3F’). The polarized distribution of ROCK activity contrasts with that of RhoA suggesting that other downstream effectors mediate its functioning during axonal elongation.

### LPA stimulates MT-stabilization in growth cones of elongating axons

In a scratch-wound migration assay with fibroblasts, LPA-RhoA-mDia signaling induces rapid polarized formation of stable MTs required for directed migration (Li and Gundersen, 2008; Bartolini and Gundersen, 2010; Etienne-Manneville, 2013; Wojnacki et al., 2014). Whether or not, this signaling pathway could induce similar changes in elongating axons remains to be established. Therefore, in the next series of experiments we first tested if LPA could alter neuronal MT organization and/or dynamics. We focused our analysis on axonal growth cones of late-stage 3 neurons, since this is the major site of MT assembly/stabilization and membrane addition required for axonal elongation (Quiroga et al., 2017). LPA (10 µM) rapidly (30 min) induced an increase in the Glu-(Fig. 4A-*B*, left panel) and acetylated α-tubulin (Supplementary Fig. 4) fluorescence signals that extend into the central and peripheral growth cone regions, usually devoid of stable MT. These changes in MT organization were exclusively found in growth cones (not observed in axonal shafts or minor neurites). The morphology of the growth cones remained unchanged; no modifications in growth cone area (Fig. 4B, middle panel) or perimeter (Fig. 4B, right panel) were detected. In accordance with these observations a significant decrease in acetylated α-tubulin immunofluorescence was also detected in axons of RhoA-suppressed neurons; this effect was quite prominent at the distal end of the axon, including the growth cone (Fig. 4C*-G*).

**Figure 4.**
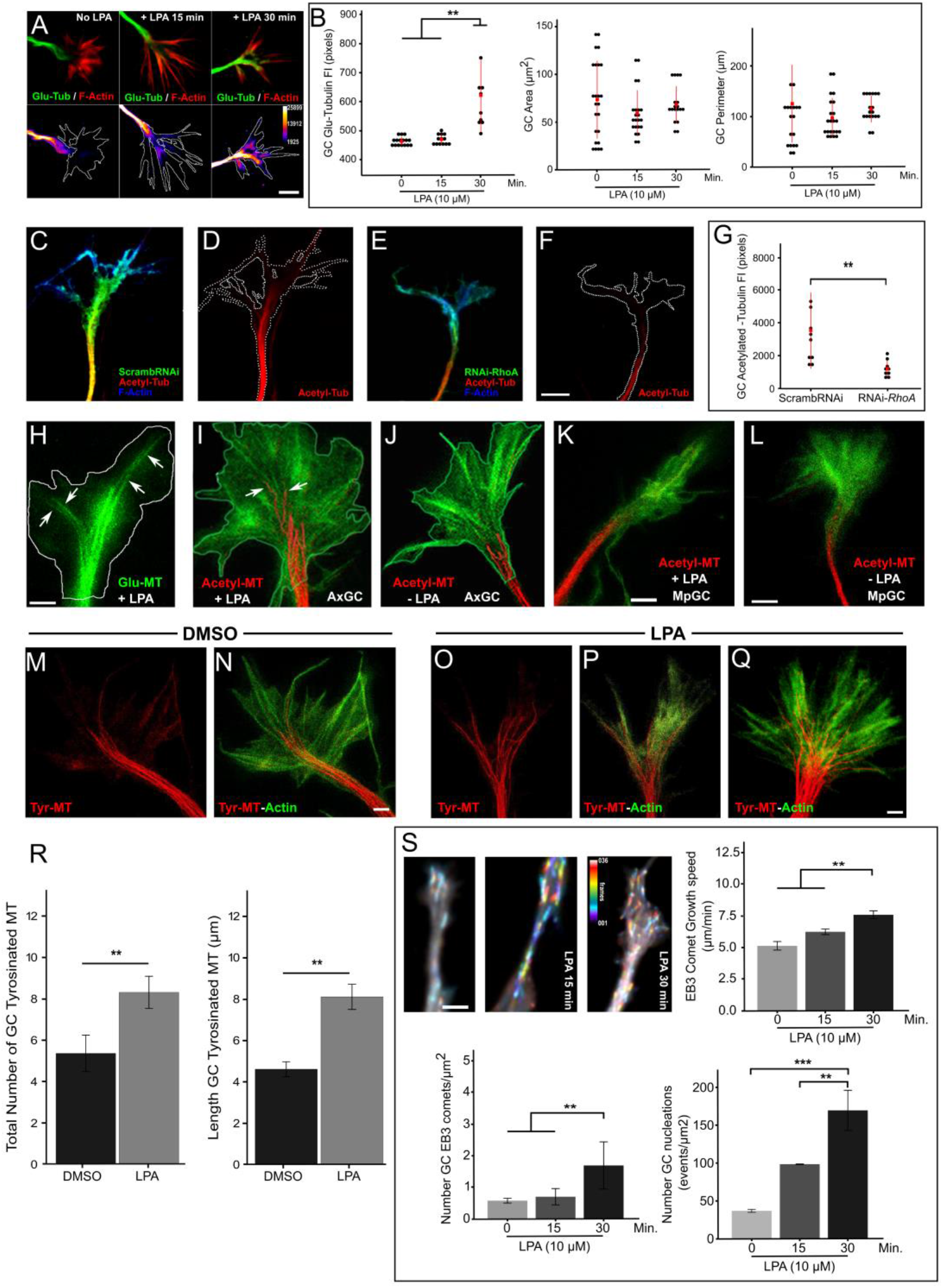
LPA promotes microtubule stabilization during axonal elongation (*A*, upper panels) Representatives confocal images of axonal growth cones of 3 DIV hippocampal neurons before and after 15 or 30 min treatments with LPA (10 µM). Fixed neurons were stained with a rabbit antibody against detyrosinated α-tubulin (Glu-tubulin, green) and phalloidin-rhodamine (red). (*A*, lower panels) Glu-tubulin channel of the axonal growth cones shown in the upper panel assigned to a pseudo-color that reflect differences in fluorescence intensity (Fire LUT, color code bar). Scale bar: 5 µm. Note the rapid increase in the Glu-tubulin fluorescence signals within growth cone area after LPA treatment. (*B*) Graphs showing the quantification of total Glu-tubulin fluorescence intensity within axonal growth cone (left panel), axonal growth cone area (middle panel) and growth cone perimeter (right panel) in control and 15-or 30-min LPA-treated neurons. Each black dot represents the value of a single growth cone. Red dots represent the mean ± S.E.M.; *<0.05; Student’s t-test. For all experiment, 18 to 24 neurons were quantified and pooled from at least three independent cultures. (*C*, *F*) Representative confocal images of axonal growth cones of 3 DIV hippocampal neurons co-transfected with scrambleRNAi + GFP (*C*, *D*) or RNAi-RhoA + GFP (*E*, *F*). Fixed neurons were stained with a mAb against Acetylated α-tubulin (red) and Phalloidin-Alexa 633 (blue). Scale bar: 5 µm. (*G*) Graph showing the quantification of Acetylated α-tubulin fluorescence intensity within axonal growth cone in *B* and *D*. Each black dot represents the value of a single growth cone. Red dots represent the mean ± S.E.M.; *<0.05; Student’s t-test. For all experiment, 9 and 10 neurons were quantified pooled from at least three independent cultures. (*H*) Representative STED image of an axonal growth cone of a 10 µM LPA-treated 3DIV neuron stained for Glu-tubulin (green). Arrows show Glu MTs extending into the central and peripheral growth cone regions. Scale bar: 2 µm. (*I*-*L*) Representatives STED images showing axonal (AxGC *I*, *J*) and minor process (MpGC *K*, *L*) growth cones from control and 10 µM LPA-treated 3DIV cultures stained with a mAb against Acetyl-tubulin (red) and phalloidin-atto594 (green). Arrows show acetylated MTs extending into the central and peripheral growth cone regions. Scale bar: 2 µm. (*M*-*Q*) Representative STED images showing axonal growth cones of DMSO (*M*, *N*) and 10 µM LPA-(*O*-*Q*) treated 3 DIV neurons stained with a mAb against tyrosinated α-tubulin (red) and phalloidin-atto594 (green). Scale bar: 2 µm. (*R*) Graphs showing quantification of total number (left panel) and length (right panel) of tyrosinated MTs profiles (see Methods) in axonal growth cones of control (DMSO-treated) and LPA (10µM)-treated 3 DIV cultures. Average of the total number of individual tyrosinated MTs (left) and average total length of tyrosinated MTs (right) in axonal growth cones are shown. Graphs represent mean ± S.E.M.; **<0.01; Student’s t-test. For all experiments, 15 neurons were analyzed and pooled from at least three independent cultures. (*S*) Representative images showing EB3-GFP comets from control neuronal cultures and treated with LPA (10 µM) for 15 or 30 min. The time lapse images are color-coded according to the time EB3 comets remain associated with MT + ends (color code scale bar). Color-code according to the scale bar. Scale bar: 5 µm. Graphs showing quantification of the mean EB3 comet growth speed (upper right), the mean number of growth cone’s EB3 comets per µm^2^ (lower left) and the mean number of growth cone’s nucleation events per µm^2^ (lower right) are shown. Graphs represent mean ± S.E.M.; **<0.01; one-way ANOVA and Tukey’s *post hoc* test. For all experiments, 10 neurons were analyzed and pooled from at least three independent cultures.

To better visualize the presence and location of Glu-or acetylated-MTs in axonal growth cones of LPA-treated neurons, we used <u>ST</u>imulated-<u>E</u>mission-<u>D</u>epletion (STED) nanoscopy. Super-resolution imaging clearly revealed more stable MTs in the central growth cone domain of LPA-treated neurons, with some of them reaching the transition and growth cone peripheral domains (Fig. 4H, I, arrows). A completely different situation was observed in axonal growth cones of control neurons, where most stable MTs only reached the base or neck of the growth cone (Fig. 4J).

The effect of LPA on stable MTs was not observed in growth cones of minor neurites of either stage-2 or stage-3 neurons (Fig. 4K*-L*).

It has been proposed that stable MT ends serve as seeds for the assembly of new MT polymers within the axonal growth cone (Mitchison and Kirschner, 1988; Witte et al., 2008; Conde and Caceres, 2009); therefore, we decided to evaluate the effect of LPA on the distribution and abundance of dynamic MTs in axonal growth cones of stage-3 hippocampal neurons. STED nanoscopy revealed that dynamic MTs, labeled with mAbs (monoclonal antibodies) that recognize tyrosinated α-tubulin, a marker of recently assembled polymer, are more abundant in LPA-treated neurons than in control ones (Fig. 4M-*Q*). Quantitative analyses of super-resolved images showed that the number and length of axonal growth cone tyrosinated MTs increased significantly after LPA treatment (Fig. 4R), despite that no changes in growth cone area were detected between LPA-and DMSO-treated cultures.

To further analyze the stimulatory effect of LPA on MT assembly we used EB3-GFP construct that binds to the plus ends of growing MTs and thus serves to monitor MT dynamics (van de Willige et al., 2016). Live imaging of growth cone EB3 comets (see Methods) revealed that it was possible to follow them for longer periods of time in LPA-treated neurons than in control ones (Fig. 4S). A quantitative analysis of trajectories of EB3 comets in axonal growth cones with the software *plusTipTracker* (Applegate et al., 2011; Stout et al., 2014) confirmed this observation, revealing that LPA treatment produced significant increments of EB3 comet speed (µm/min), of the mean number of EB3 comets per/unit area, and of the mean number of nucleation events per/unit area (Fig. 4S). These observations are fully consistent with those derived from the immunofluorescence data obtained using either confocal microscopy or nanoscopy (Fig. 4A*-R*). Importantly, no stimulatory effect of LPA on MT abundance was detected in growth cones of minor neurites; on the contrary, they appear to contain less MTs, and fewer ones penetrating into growth cones (Supplementary Fig. 5).

### Formin are required for axon elongation

Formins are another type of RhoA effectors that regulate the actin and MT cytoskeletons (Goode and Eck, 2007; Bartolini and Gundersen, 2010; Kuhn and Geyer, 2014); some of them, like Mammalian Diaphanous (mDia), have been implicated in neuronal tangential migration (Shinohara et al., 2012), axonal guidance (Toyoda et al., 2013), dendrite complexity and spine density in differentiated hippocampal neurons (Qu et al., 2017), as well as axonal regeneration (Pinto-Costa et al., 2020) and SDF-1 α-promoted axonal elongation in cerebellar neurons (Arakawa et al., 2003). Since we did not detect any effect on axonal elongation upon ROCK inhibition, it became of interest to evaluate if the RhoA stimulatory effect on axon elongation involves formins. To this end, we used the <u>s</u>mall <u>m</u>olecule inhibitor of formin <u>h</u>omology <u>2</u> domains (SMIFH2) that effectively inhibits formin-mediated actin nucleation (Rizvi et al., 2009) and MT assembly (Isogai et al., 2015; Qu et al., 2017). Addition of SMIFH2 to freshly plated hippocampal neurons did not revert the inhibitory effect of LPA on axon outgrowth (Fig. 5A) suggesting that RhoA-formin is not part of the signaling pathway inhibiting neurites from engaging in axon-like growth. However, SMIFH2 alone did cause a 15% decrease in the number of neurons progressing to stage 3 (Fig. 5A), suggesting that at least one formin might be involved in axon outgrowth, acting as a positive growth-promoting signal rather than as a negative one. A similar experiment (a 24 h treatment with SMIFH2) carried out after 3 DIV, when almost all neurons already extended an axon, revealed that SMIFH2 treatment significantly decreases axonal length and LPA-enhanced axonal elongation (Fig. 5B, left panel), without altering the number or length of minor neurites (Fig. 5B, right panel). Together, these results suggest that formins exerts a weak stimulatory and strong inhibitory influence on axon outgrowth, but only a stimulatory one during elongation. The growth-promoting activity of RhoA on axon growth appears to be mediated by a formin. We further tested this possibility through experiments of axon regeneration.

**Figure 5.**
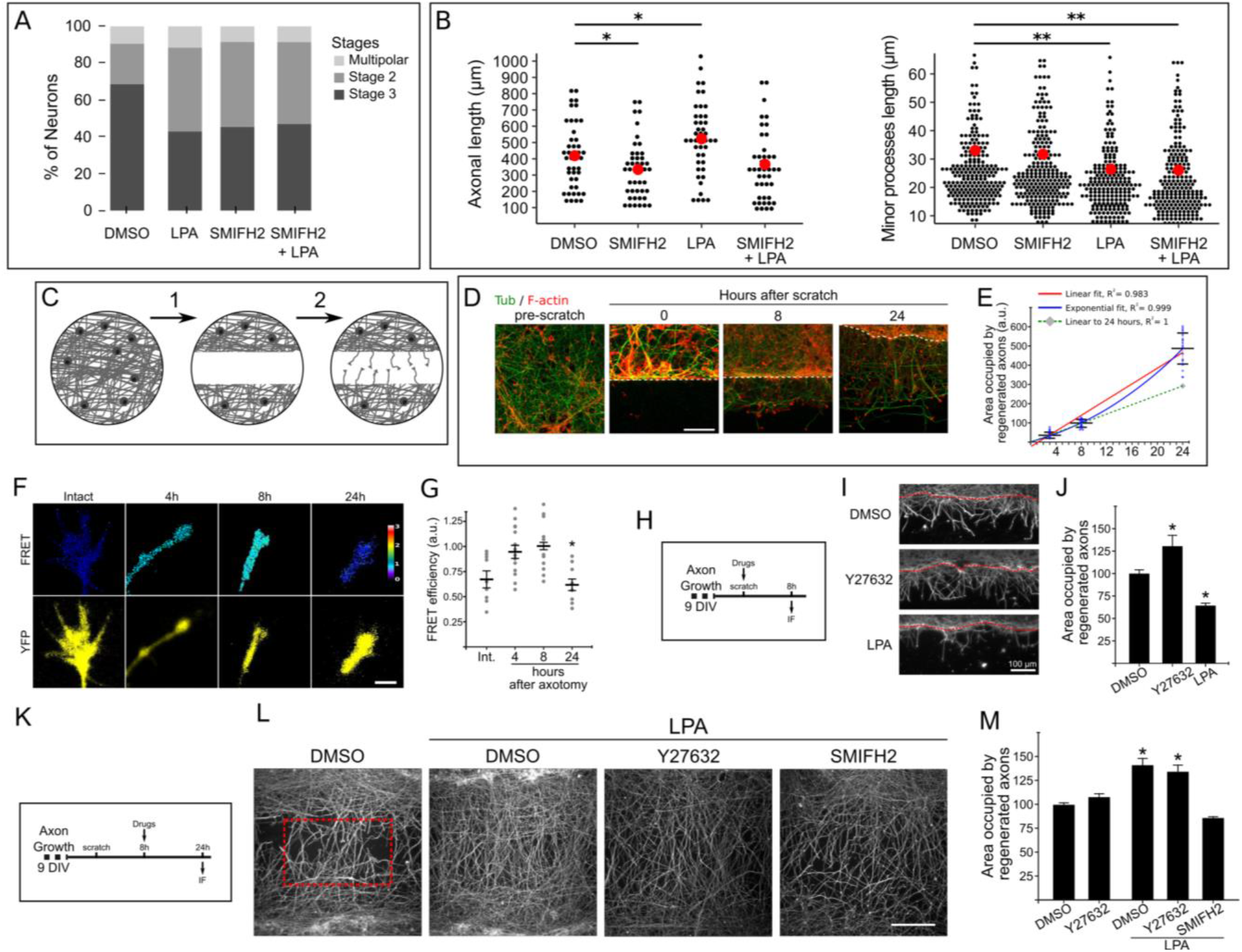
LPA stimulated axonal elongation is mediated by formins. *(A)* Graph showing the percentage of stage2, stage 3 and multipolar neurons in cultured hippocampal neurons treated for 36 h with vehicle (DMSO), LPA (10 µM), SMIFH2 (10 µM) and SMIFH2 + LPA (10 µM + 10 µM). Percentages were calculated from 3 independent cultures. (*B*) Graphs showing quantifications of the total axonal (left panel) and minor processes (right panel) lengths of 3DIV neurons treated for 24 h with vehicle (DMSO), SMIFH2 (5 µM), LPA (10 µM) or SMIFH2 + LPA (5 µM + 10 µM). Each black dot represents the neurite length of a single neuron. Red dots represent the mean neurite length ± S.E.M; *< 0.05, **< 0.01; one-way ANOVA and Tukey’s *post hoc* test. For all experiment, 35 to 44 neurons were quantified and pooled from at least three independent cultures. (*C*) Schematic representation of the cortical scratch assay used to evaluate axon regeneration in cultured cortical neurons. Primary cortical neurons growth for 9 DIV were mechanically injured and cleared by a scratch (1); axonal re-growth into the scratched area (i.e. regeneration, (2)) was then evaluated 4, 8 and 24 h later. (*D*) Representative confocal images of cortical neuronal before and after different periods following the scratch. The cultures were stained with a mAb against tyrosinated α-tubulin (green) and rhodamine-phalloidin (red). The dashed line indicates the border of the scratched area. By 8 h after axotomy, regenerating axons began to populate the scratch showing F-actin-rich growth cones. Scale bar: 50 µm. (*E*) Graph showing the quantification of the area occupied by regenerated axons at 4, 8 and 24 h after the scratch. Different curves were fitted to the observed values to visualize possible changes in growth rate. Note that the linear fit showed the lowest regression value (red line), while the exponential curve shows the best fit (blue line) indicating that the growth rate is low within the first 8 h, but that after that period increases over time. The linear fit including the early time points (dashed green line) shows that the expected value at 24 h if growth rate was constant (grey diamond) is more than 2 standard deviations away from the observed growth. Graphs represent mean ± S.E.M. 18 areas (ROI) occupied by regenerated axons were quantified and pooled from three independent cultures. (*F*) Representative FRET map images showing ROCK activity in 9 DIV intact and regenerating axonal growth cones at 4, 8 and 24 h after the scratch-induced axotomy. FRET measurements were performed using the unimolecular Eevee-ROCK FRET-based biosensor. A ratiometric method was used to measure the FRET signal and the FRET maps are color-coded according to activation intensity (color-code scale bar). Scale bar: 5 µm. (*G*) Graph showing quantification of ROCK activity in intact and regenerated axonal growth cone. Each small dot represents the FRET value in a single growth cone. Each large red dot represents the mean ± S.E.M.; *< 0.05; one-way ANOVA and Tukey’s *post hoc* test. 9 to 13 regenerated growth cones were quantified and pooled from three independent cultures. (*H*) Schematic representation of the experimental design to evaluate the LPA-RhoA pathways during axon re-growth after 8 h of the scratch lesion. (*I*) Representative images of regenerated axons after 8 h of axotomy in the continuous presence of vehicle (DMSO), 10 µM Y27632 or 10 µM LPA. Scale bar: 100 µm. (*J*) Graph showing the quantification of (*I*). The area occupied by regenerated axons into the scratch normalized to 100 in the control condition is shown. Graphs represent mean ± S.E.M.; *< 0.05; one-way ANOVA and Tukey’s *post hoc* test. 14 to 20 areas were quantified and pooled from three independent cultures. (*K*) Schematic representation of the experimental design to evaluate the LPA-RhoA pathways during axon re-growth after 24 h of the scratch lesion. (*L*) Representative images of control regenerated axons after 24 h of axotomy and treated with LPA alone or LPA + Y27632 or LPA + SMIFH2; all drugs were added 8 h after axotomy. The dotted (red) rectangle delimits the area where regenerated axons were quantified. Scale bar: 200 µm. (*M*) Graph showing the quantification of the area occupied by regenerated axons into the scratch in the presence of vehicle (DMSO), Y27632 (10 µM), LPA (10 µM), LPA + Y27632 (10 µM + 10 µM) and LPA + SMIFH2 (10 µM + 10 µM). Normalized to 100 in the control condition (vehicle). Graphs represent mean ± S.E.M.; *< 0.05; one-way ANOVA and Tukey’s *post hoc* test. 10 to 36 areas were quantified and pooled from three independent cultures.

It is now generally accepted that nerve injury activates a RhoA-ROCK signaling pathway leading to inhibition of axon regeneration (McKerracher et al., 2012; Fujita and Yamashita, 2014). With the exception of a role ascribed to formins in mediating the effect of profilin1 in promoting MT growth in growth cones (Pinto-Costa et al., 2020), the involvement of formins in this setting has remained largely unexplored. Therefore, we performed a series of experiments to test if an LPA-RhoA-formin pathway could stimulate axon growth after injury. To this end, we used a cortical scratch assay, where rat cortical neurons are grown to allow extensive axon elongation for 9 days and then mechanically injured with a pipet tip (Fig. 5C*-D*; (Huebner et al., 2011; Kaplan et al., 2017)); one day later, regenerating axon-like neurites (Tau-1 +) invade the scratched area.

In the first set of experiments, we evaluated axon regeneration by quantifying the area occupied by neurites in the scratch region at different time points after the lesion (Fig. 5E). This analysis revealed that regenerative axon growth proceeds slowly during the first 8 h post-transection, but afterwards accelerates displaying an exponential rate (Fig. 5E). RhoA-ROCK activity was also evaluated at the same time points after axon transection. FRET imaging showed that both RhoA (not shown) and ROCK (Fig. 5F*-G*) activities increased significantly shortly (4, 8 h) after axon injury. At later time points (24 h), RhoA activity remains high while that of ROCK decreases (Fig. 5F*-G*), showing lower levels in fast extending axons.

In the next series of experiments, we applied Y27632 or LPA at the time of axon transection, and quantified regeneration 8 h later (Fig. 5H*-J*). We found that inhibition of ROCK with Y27632 significantly enhanced axon regeneration (Fig. 5I-*J*) while LPA had the opposite effect (Fig. 5I-*J*). To evaluate the possible existence of a growth-promoting effect of RhoA on axon regeneration, LPA was applied 8 h after injury (Fig. 5K-*M*). In this setting, LPA significantly increased regeneration, which was blocked by SMIFH2, but not by Y27632 (Fig. 5L, M), suggesting that this stimulatory effect of LPA could be mediated by a formin. The results on regenerating axons recapitulate our finding in developing neurons: RhoA has ROCK-dependent inhibitory role in the early phases, and a formin-dependent stimulatory role in the later phases of axonal re-growth.

### mDia promotes axonal elongation by stabilizing axonal MTs

As SMIFH2 inhibits all formins having an FH2 domain, in the final set of experiments we decided to specifically address mDia participation in axon elongation. mDia formins have a prominent role in stabilizing MTs in nonneuronal (Palazzo et al., 2001; Wen et al., 2004; Bartolini et al., 2008) and neuronal cells (Qu et al., 2017), but have not been implicated in axon specification or elongation. We silenced mDia1 by transfecting neurons with a specific shRNAi targeting mDia1 (Qu et al., 2017; see Methods) 24 hours after plating; 48 hours later we detected a substantial reduction in axonal length when compared to control shScramble-transfected neurons. Notably, the length of minor processes remained relatively unaffected (Fig. 6A-*C*).

**Figure 6.**
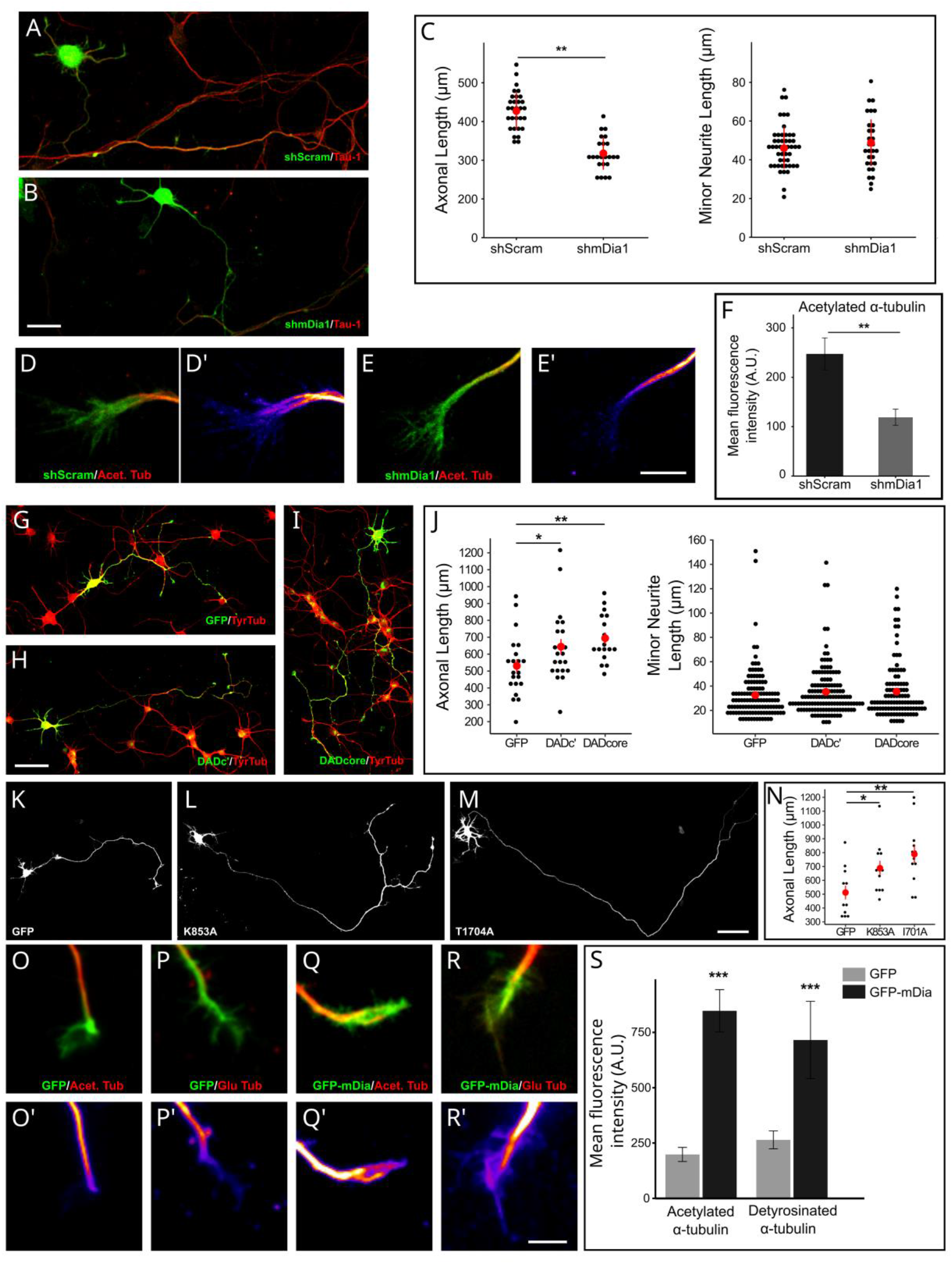
RhoA effector mDia1 promotes axonal elongation. (*A, B*) Representative confocal images of cultured hippocampal neurons (3 DIV) transfected with scrambled control (shScram; *A*) or shRNA against mDia1 (shmDia1; *B*) stained with anti-Tau-1 mAb (red). Cultures were transfected 12 h after plating an examined at 3DIV. Scale bar: 10 µm. (*C*) Graphs showing the quantification of total axonal (left) and minor processes length (right) in neurons expressing control shScram or shmDia1. Each black dot represents the neurite (axon or minor process) length of a single neuron. Red dots represent the mean neurite length ± S.E.M; **<0.01; Student’s t-test. For all experiment, 25 to 30 neurons were quantified and pooled from at least three independent cultures. (*D, E*) Representative confocal images showing growth cones of 3 DIV cultured hippocampal neurons transfected with control shScram (green; *D*) or shmDia1 (green; *E*) and stained with the mAb against Acetylated-tubulin (red; *D*, *E*). (*D*’, *E’*) Acetylated-tubulin channel of the axonal growth cones shown in (*D, E*) assigned to a pseudo-color scale that reflect differences in pixel value (Fire LUT, color code bar). Scale bar: 5 µm. Note the decrease in the Acetylated tubulin fluorescence signals within growth cone area in neurons expressing shmDia1. (*F*) Graph showing quantification of total mean Acetylated tubulin fluorescence intensities within axonal growth cones of control shScram or shmDia1 transfected neurons. Graphs represent mean ± S.E.M.; **<0.01; Student’s t-test. For all experiment, 20 neurons were quantified and pooled from at least three independent cultures. (*G-I*) Representative confocal images of cultured hippocampal neurons (3 DIV) transfected with GFP (*G*), GFP mDia DADc’ domain (*H*) and GFP mDia DADcore domain (*I*) stained with anti-tyrosinated tubulin mAb (red). Cultures were transfected 60 h after plating an examined 18 h later. Scale bar: 50 µm. (*J*) Graphs showing the quantification of total axonal (left) and minor processes length (right) in neurons expressing control GFP, mDia DADc’ or mDia DADcore domains. Each black dot represents the neurite (axon or minor process) length of a single neuron. Red dots represent the mean neurite length ± S.E.M; *<0.05; **<0.01; one-way ANOVA and Tukey’s *post hoc* test. For all experiment, 17 to 21 neurons were quantified and pooled from at least three independent cultures. (*K-M*) Representatives confocal images of cultured hippocampal neurons (3 DIV) transfected with eGFP (*K*), mDia mutant K853A (*L*) and mDia mutant T1704A (*M*). Scale bar: 50 µm. (*N*) Graph showing the quantification of total axonal length in neurons expressing GFP or mDia mutants. Each black dot represents the total axonal length of a single neuron. Red dots represent the mean total axonal length ± S.E.M.; *<0.05; **<0.01; one-way ANOVA and Tukey’s *post hoc* test. For each condition, 12 neurons were quantified from three independent experiments. (*O-R*) Representative confocal images showing growth cones of 3 DIV cultured hippocampal neurons transfected with control GFP (green; *O*, *P*) or GFP-mDia (green; *Q*, *R*) and stained with the mAb against Acetylated-tubulin (red; *O*, *Q*) or the rabbit polyclonal antibody against Glu-tubulin (red; *P*, *R*). (*O*’-*R’*) Acetylated (*O*’, *Q’*) and Glu (*P’, R’*) -tubulin channel of the axonal growth cones shown in (*O-R*) assigned to a pseudo-color scale that reflect differences in pixel value (Fire LUT, color code bar). Scale bar: 5 µm. Note the increase in the Glu and Acetylated-tubulin fluorescence signals within growth cone area in neurons expressing GFP-mDia. (*S*) Graph showing quantification of total mean Acetylated and detyrosinated (Glu-) tubulin fluorescence intensities within axonal growth cones of GFP and GFP-mDia transfected neurons. Graphs represent mean ± S.E.M.; ***<0.01; Student’s t-test. For all experiment, 35 neurons were quantified and pooled from at least three independent cultures.

Consistent with the increase in stable MTs after LPA treatment (Figure 4), quantitative analysis revealed that neurons expressing shmDia1 exhibited a lower acetylated α-tubulin fluorescence signal compared to control neurons (Fig. 6D-*F*). These results suggest that mDia1 is necessary for microtubule polymerization and stabilization within the growth cones, consequently facilitating axonal elongation.

Diaphanous-related formins are auto inhibited through intramolecular binding of a <u>d</u>iaphanous <u>a</u>utoregulatory <u>d</u>omain (DAD) to a conserved N-terminal <u>d</u>iaphanous inhibitory <u>d</u>omain (DID) (Rose et al., 2005). Binding of active RhoA to DID displaces DAD from the N-terminal region, releasing auto-inhibition and therefore activating mDia1 (Lammers et al., 2005; Otomo et al., 2005). Transfection with a GFP-tagged mDia1 DAD domain fragment activates the endogenous protein by relieving auto-inhibition (Alberts, 2001; Palazzo et al., 2001; Eng et al., 2006). Two different GFP-tagged mDia1 DAD domains, C’ terminal and core domains (Alberts, 2001; Wallar et al., 2006), were used to explore the effect of endogenous mDia activation on axonal length. To this end, axonal length was quantified in cultured hippocampal neurons transfected with GFP or GFP-DAD constructs (GFP-DAD core or GFP-DAD-Ć) at 2 DIV and evaluated 24 h later. DAD-mediated mDia activation significantly increased axonal length when compared to control (GFP-transfected) cultures (Fig. 6H-*I*, 6*J*, left panel). By contrast, and consistent with SMIFH2 effects, ectopic expression of either GFP-DAD core or GFP-DAD-Ć did not affect minor neurite length (Fig. 6J, right panel). Together, these results point towards mDia being the formin activated downstream of RhoA during axonal elongation.

mDia is best known for stimulating actin nucleation and polymerization (Pruyne et al., 2002). However, in recent years it has become increasingly evident that it also promotes MT stabilization in non-neuronal cells and in primary neurons (Palazzo et al., 2001; Bartolini et al., 2008; Bartolini and Gundersen, 2010; Bartolini et al., 2012; Qu et al., 2017). The MT-stabilizing domain of mDia2 maps within the same region involved in actin polymerization, namely the FH2 domain (Xu et al., 2004). Two different point mutations (K853A and I704A) within the FH2 domain of constitutive active mDia prevent actin polymerization activity but retain the MT-stabilizing activity (Harris et al., 2006; Bartolini et al., 2008). Because an LPA-RhoA-mDia MT-stabilization signaling pathway is required for polarized cell migration in fibroblasts (Bartolini et al., 2008; Morris et al., 2014), and because LPA-promoted-axonal elongation also involves enhanced MT stabilization it became of interest to test the consequences of expressing theses mutants on axon extension. To this end, 2 DIV hippocampal cell cultures were transfected with either K853A or I704A mDia mutants or a control plasmid and axonal length evaluated 18 h later. Ectopic expression of any one of these mutants resulted in a significant increase in axonal length (Fig. 6K-*N*). It is noteworthy that this effect was similar to the one observed after endogenous mDia activation induced by transfection of DAD domains (Fig. 6G*-J*).

Then, we addressed the question of whether mDia could promote axonal elongation by modifying the dynamic behavior of MTs. To test this idea 2 DIV hippocampal neurons were transfected with GFP-mDia (wild type) or the I704A mDia mutant and 1 day later changes in the relative amount of stable MTs were evaluated by quantitative immunofluorescence with antibodies against Glu-or acetylated-α-tubulin. The results obtained revealed that the ectopic expression of either mDia (Fig. 6O-*R*) or the MT-stabilizing mutant I704A (not shown) significantly increase the fluorescent signal generated by both antibodies (Fig. 6S). Together, this information supports the possibility that mDia-promoted MT-stabilization mediates RhoA-enhanced axonal elongation.

## Discussion

### Dual functions of RhoA in axon growth

It is now well established that small Rho-GTPases play major roles in neuronal morphogenesis by controlling cytoskeletal organization/dynamics and membrane trafficking (Gonzalez-Billault et al., 2012; Wojnacki and Galli, 2016; Quiroga et al., 2017). A long-lasting view based on pharmacological studies and evaluation of the up-or down-regulation of their expression/activity during neuronal development, has established the idea of RhoA being a major inhibitory factor for the novo and regenerative axon growth (Arimura and Kaibuchi, 2007; McKerracher et al., 2012; Schelski and Bradke, 2017; Takano et al., 2017; Dupraz et al., 2019; Takano et al., 2019; Wilson et al., 2020b). The present results put forward a far more complete picture of the function of RhoA during axon formation and growth. First, FRET imaging assays conclusively challenge the current dogma by revealing First, FRET imaging assays conclusively challenge the current dogma, by revealing not only high RhoA activity in undifferentiated neurites exhibiting little if any growth, but also and surprisingly, in elongating axons. A similar behaviour was observed after axotomy, where high RhoA activity was detected in both slow and fast regenerating axons. Second, along with RNAi and pharmacological experiments, our observations demonstrate that RhoA can function as a positive or negative regulator of axon growth depending on various factors such as neurite type (minor neurite vs. prospective axon vs. elongating axon), stage of development (axon outgrowth vs. axon elongation), context (slow regenerating vs. fast regenerating axons) and downstream effectors (e. g. ROCK vs. mDia; see next section). Moreover, we reveal for the first time that dual inhibitory and stimulatory functions of RhoA co-exist at the same time within the same cell but at different domains, as in the case of late stage-3 cultured hippocampal pyramidal neurons, where active RhoA promotes axonal elongation while preventing minor neurite growth and the subsequent generation of multiple axon-like neurites.

All this information is consistent with data from another important morphogenetic event, namely cell migration, where Rho GTPases also serve as master regulators (Etienne-Manneville, 2013). In this setting, early studies prompted the idea of Rac/Cdc42 acting as stimulatory factors promoting actin protrusive activity and cell locomotion at the leading edge, and RhoA being a negative regulator triggering acto-myosin contractility and cell retraction at the trailing edge (Raftopoulou and Hall, 2004). However, it soon became evident that this impression was an oversimplification and that depending on cell type and environmental factors, considerable variations exist in the spatio-temporal activation pattern and function of RhoA (Kurokawa and Matsuda, 2005). For example, in migrating HeLa cells, FRET experiments revealed RhoA activation at both the leading and trailing edges, while in migrating MDCK cells RhoA activity was only detected at the leading edge (Kurokawa and Matsuda, 2005); interestingly, in growth factor-stimulated migrating fibroblasts RhoA activity was low at the plasma membrane, but high at membrane ruffles of growing lamellipodia, where high Rac activity was also detected (Kurokawa and Matsuda, 2005). Subsequent studies with a new generation of Rho-GTPase FRET biosensors and computational multiplexing revealed that RhoA activation at the leading edge of migrating fibroblasts is synchronized with protrusion and precedes Rac-Cdc42 activation (Pertz et al., 2006; Machacek et al., 2009). Furthermore, it was proposed that this early activation of RhoA might serve to initiate formin-mediated actin polymerization. Altogether, this information clearly indicates that in other cell types, RhoA has both stimulatory and inhibitory activities spatially confined to particular cellular domains, being both required for proper cell growth and polarization.

### Opposite functions of ROCK and mDia on axon growth

It has been proposed that different subcellular pools of a given Rho-GTPase associated with distinct downstream effectors and regulatory proteins comprise a “*spatio temporal signaling module*” (Pertz, 2010) that performs different functions when associated with distinct downstream effectors and regulatory proteins. Our experiments show that in cultured hippocampal neurons, the inhibitory effect of RhoA on axon formation involves ROCK while a different effector, namely mDia, has an opposite influence stimulating axon elongation.

Previous pharmacological studies have established that inhibition of ROCK in early developing cultured hippocampal neurons promotes axon formation and/or the generation of multiple axon-like neurites (Takano et al., 2019); see also References in the Introduction of this study). More recently, an optogenetic approach showed that activation of RhoA or ROCK at the cell body or distal end of newly formed axons results in retraction of minor neurites or inhibition of axon outgrowth, respectively (Takano et al., 2017). Consistent with this, our FRET experiments revealed high activity of RhoA and ROCK in growth cones of minor neurites and a simultaneous decrease of activity selectively located at the large growth cones of prospective and newly formed axons. However, while ROCK activity decreases significantly and permanently from elongating axons, RhoA activity show a second rise after axon sprouting becoming high in axonal shafts and growth cones of late stage-3 neurons. A similar dissociation of RhoA and ROCK activities was observed after axotomy in cultured cortical neurons. Slow and fast regenerative axon growth correlates with high and low ROCK activities, respectively, but not with equivalent changes in RhoA activity. As in the case of cultured hippocampal neurons, pharmacological experiments support the idea of RhoA-ROCK acting as a growth inhibitory signal while RhoA-formin as a stimulatory one.

Activation of endogenous mDia by transfection of cultured hippocampal neurons with a GFP-tagged <u>d</u>iaphanous <u>a</u>uto inhibitory <u>d</u>omain (DAD) reveals that mDia mediates the stimulatory effect of RhoA on axonal elongation, while its down-regulation with a specific shRNAi have the opposite effect. These results are consistent with a previous report showing that in cultured cerebellar granule cells stromal-cell derived factor (SDF-1α) promotes axonal elongation by activating a RhoA-mDia signaling pathway (Arakawa et al., 2003). We have now extended this observation by showing not only that RhoA-mDia stimulates axonal elongation in the absence of exogenous SDF-1α, but also by providing evidence about its mechanism of action.

Our results indicate that RhoA-mDia signaling regulates axonal extension this effect by promoting MT-stability and assembly at growth cones of elongating axons. Several lines of evidence support this notion. First, using different experimental approaches we showed that LPA increases the number of stable MTs entering the central and peripheral domains of axonal growth cones. Second, this effect was prevented by SMIFH2 and shmDia1, unaffected by Y27632 and absent in growth cones of minor neurites. Third, mDia mutants lacking actin polymerization activity, but retaining their ability to promote MT stabilization stimulate axonal elongation. Finally, these observations are in full agreement with studies in scratched monolayers of NIH-3T3 fibroblasts showing that LPA-induced RhoA activation drives mDia mediated-MT stabilization at the leading edge required for polarized cell migration (Cook et al., 1998; Palazzo et al., 2001; Gundersen et al., 2005; Bartolini and Gundersen, 2010; Bartolini et al., 2012).

The precise mechanism by which mDia stimulates MT stabilization in elongating axons remains to be established. In this regard, it is worth noting that KIF4, a kinesin superfamily member highly enriched in growth cones of growing axons and required for L1-stimulated axonal elongation (Peretti et al., 2000), interacts with EB1 to mediate mDia-induced MT stabilization in migrating fibroblasts (Morris et al., 2014). Future studies should explore this, among others possibilities.

## Methods

### Animal use and care

Pregnant Wistar rats were born in the vivarium of INIMEC-CONICET-UNC (Córdoba, Argentina). Wistar rats lines were originally provided by Charles River Laboratories International Inc (Wilmington, USA). All procedures and experiments involving animals were approved by the Animal Care and Ethics Committee (CICUAL http://www.institutoferreyra.org/en/cicual-2/) of INIMEC-CONICET-UNC (Resolution numbers 014/2017 B, 015/2017 B, 006/2017 A and 012/2017 A) and were in compliance with approved protocols of the National Institute of Health Guide for the Care and Use of Laboratory Animals (SENASA, Argentina).

All methods were carried out in accordance with relevant guidelines and regulations.

### DNA Constructs and siRNAs

The following cDNA constructs were used in this study: 1) cDNA coding for small RhoGTPases biosensors Rac1-Raichu (Itoh et al., 2002) (Gently provided by Dr. Matsuda, Institute of Advanced Energy, Kyoto University, Japan); RhoA 1G and RhoA 2G (Fritz et al., 2013) (Gently provided by Dr. Pertz, Institute of Cell Biology, Bern, Switzerland) 2) cDNA coding for a FRET reporter for ROCK activity designated as EeveeROCK (Li et al., 2017) (Gently provided by Dr. Matsuda, Institute of Advanced Energy, Kyoto University, Japan) 3) cDNA coding for EB3-GFP (Gently provided by Dr. Gundersen, Department of Pathology, Anatomy and Cell Biology, Columbia University, New York, USA). 4) cDNA codings for mDia1-GFP, the point mutated K853A mDia-GFP, the point mutated T1704A mDia-GFP, the DADc’ mDia-GFP domain and the DAD core mDia-GFP domain (Bartolini et al., 2008) (Gently provided by Dr. Gundersen, Department of Pathology, Anatomy and Cell Biology, Columbia University, New York, USA). 5) cDNA coding eGFP protein (Clontech). 6) Short hairpins RNA (shRNA) against mDia1 and a scrambled sequence as described and validated previously by Qu et al (Qu et al., 2017) targeting the following sequence of rat mDia1 5’ gcgacggcggcaaacataa 3’ and control scramble 5’ ggcaaatcttctagtctat 3’ . (Gently provided by Dr. Bartolini, Department of Pathology, Anatomy and Cell Biology, Columbia University, New York, USA). 7) For RhoA gene silencing experiments, siRNA oligos and RNAi negative control were purchased from GenBiotech as described and validated previously by Dupraz et al (Dupraz et al., 2019), targeting the following sequence of rat RhoA: 5’ tagtttagaaaacatcccaga 3’.

### Cell cultures

Embryonic day-18 rat embryos (euthanized by CO2 overdose) were used to prepare primary hippocampal cultures as previously described (Bisbal et al., 2018; Wilson et al., 2020a). Briefly, hippocampi from 18 days fetal rats were dissected and incubated with trypsin (0.25% for 15 min at 37°C) (Thermo Fisher Gibco; Cat Number: 15090-046) and mechanically dissociated by trituration with a Pasteur pipette. Cells were plated on Cover Glasses, Circles, 12 mm (Marienfeld Superior; Cat Number: 633029) coated with 1 mg/mL poly-L-lysine (Sigma Chemical Co; Cat Number: P2636) at a density of 2000 cells/cm2 in minimum essential medium (MEM, Thermo Fisher Gibco; Cat Number: 61100-061) supplemented with CTS GlutaMAX I Supplement (Thermo Fisher Gibco; Cat Number: A1286001), Sodium Piruvate (Thermo Fisher Gibco; Cat Number: 11360070), Penicillin-Streptomycin (Thermo Fisher Gibco; Cat Number: 15140122), and 10% horse serum (Thermo Fisher Gibco; Cat Number:16050122). After 2 h, the coverslips were transferred to dishes containing serum-free Neurobasal (Thermo Fisher Gibco; Cat Number: 21103049) with B-27 Plus Supplement (Thermo Fisher Gibco; Cat Number: A3582801) and CTS GlutaMAX I Supplement (Thermo Fisher Gibco; Cat Number: A1286001). For cortical scratch assays, cortical neurons were prepared as described by (Kaplan et al., 2017) plated on 12 mm cell culture coverslips at a density of 200000 cells/cm2 using same neuronal plating and maintenance media than the used for hippocampal cultures.

CHO cells (ATCC) were plated on round 12-mm cover glasses (Marienfeld Superior; Cat Number: 633029) and cultured in Minimum Essential Medium (MEM, Thermo Fisher Gibco; Cat Number: 61100-061) supplemented with 10% fetal bovine serum (Internegocios S.A., Cat Number: 000008), 1% CTS GlutaMAX I Supplement (Thermo Fisher Gibco; Cat Number: A1286001), 1% sodium pyruvate (Thermo Fisher Gibco; Cat Number: 11360070) and 1% penicillin streptomycin (Thermo Fisher Gibco; Cat Number: 15140122) at 37°C, 5% CO2, until fixation.

### Cell transfection and immunofluorescence

Transient electroporation: Cultured neurons were electroporated using the Lonza Nucleofector II device (Cat Number: AAD-1001N, Lonza; program O-003) and the mouse neuron Nucleofector® Kit (Cat Number: VPG-1003, Lonza), according to the manufacturer’s instructions. 0.5-1 x 106 cells were used for each transfection with 3 µg of plasmid DNA. For the silencing experiments, 0.5-1 x 106 cells were transfected with a mix of 0.2 pmol of the siRNA and 1 µg of GFP expressing plasmids (pEGFP) as carrier DNA and transfection reporter.

Transient Lipofection: Cultured cells were transfected with Lipofectamine 2000 (Thermo Fisher Gibco; Cat Number: 11668027), following manufacturer’s instructions. Briefly, for a 35 mm dish 4 μl of Lipofectamine 2000 and selected plasmids (3 μg) were diluted in Opti-MEM medium (Thermo Fisher Scientific) to a final volume of 250 μl and incubated for 20 min at RT. Cells were incubated for 2 h in Opti-MEM containing the Lipofectamine/plasmids mix and returned to Neurobasal-B-27 plus medium until analysis. For the silencing experiments, neurons were transfected with a mix of 0.2 pmol of the siRNA and 1 µg of GFP expressing plasmids (pEGFP) as carrier DNA and transfection reporter.

### Immunocytochemistry

Cells were fixed with 4% paraformaldehyde (Sigma-Aldrich, Cat Number: 441244) and 4% sucrose diluted in phosphate buffered saline (PBS) for 20 min at RT as previously described (Bisbal et al., 2018; Pesaola et al., 2021). For STED and confocal growth cones MTs analysis neurons were simultaneously fixed and permeabilized in PHEM buffer (60 mM PIPES, 25 mM HEPES, 5 mM EGTA, 1 mM MgCl (pH 6.9)) containing 0.25% glutaraldehyde (Sigma-Aldrich, Cat Number: G6257), 3.7% paraformaldehyde, 3.7% sucrose and 0.1% Triton X-100, and quenched in 0.1M Glycine/PBS for 10 min as previously described (Unsain et al., 2018).

Fixed cells were washed 3 times with PBS, permeabilized in 0.2% Triton X-100 in PBS at RT for 5 min and again washed in PBS before antibody incubation. Cells were incubated in blocking buffer (bovine serum albumin 5%/PBS) for 1 h. Cells were then incubated with primary antibodies diluted in blocking buffer for 1 h, washed with PBS three times and incubated with fluorescent-conjugated secondary antibodies for 1 h. Finally, cells were washed with PBS three times and the coverslips mounted using FluorSave (Millipore Calbiochem; Cat Number: 34-578-9).

### Antibodies and Reagents

The following primary antibodies were used for immunofluorescence (IF) in this study: a monoclonal antibody (mAb) against tau protein (clone Tau-1; Millipore, Cat Number: MAB3420, RRID: AB_94855) diluted 1:1000; a rat monoclonal against tyrosinated α-tubulin (clone YL1/2, Abcam, Cat. Number: ab6160, RRID: AB_305328) diluted 1:2000; a mAb against tyrosinated α-tubulin (clone TUB-1A2; Sigma, Cat. Number: T9028, RRID: AB_261811) diluted 1:1000 or 1:200 for STED microscopy; a mAb against α-tubulin (clone α-3A1; Sigma-Aldrich, Cat. Number T5168, RRID:AB_477579) dilute 1:1000; a mAb against Acetylated α-tubulin (clone 6-11B-1; Sigma-Aldrich, Cat. Number: T7451, RRID: AB_609894) diluted 1:1000 or 1:200 for STED microscopy and a rabbit polyclonal antibody against detyrosinated (Glu) tubulin (gently provided by Dr. Greg Gundersen) diluted 1:1000 or 1:100 for STED microscopy.

The following secondary antibodies and dyes were used: for widefield the corresponding secondary antibodies Alexa Fluor 488, Alexa Fluor 568 (Thermo Fisher Scientific, Cat Numbers: A11001, A11008, 11006, A11004, A11031 and A11077), phalloidin-Rhodamine and phalloidin-Alexa Fluor 647 (Thermo Fisher Scientific, Cat Numbers: R415 and A22287); and for STED microscopy anti-mouse Atto 647N, anti-rabbit Atto 594 (Sigma-Aldrich, Cat Numbers: 50185 and 77671) and phalloidin-Atto 594 (Sigma-Aldrich, Cat Numbers: 51927) was used.

LPA and SMIFH2 were purchased from Sigma-Aldrich (Cat Numbers: L7260 and S4826), Y27632 from Calbiochem (Cat Number: 688000) and C3 toxin (Rho Inhibitor I) from Cytoskeleton (Cat Number: CT04).

### Confocal microscopy

Cells were visualized using either a conventional LSM 800 (Zeiss, Germany) and TCS FV300 (Olympus, Japan) or a spectral FV1200 (Olympus, Japan) inverted confocal microscope. For high-magnification analysis, z stack confocal imaging was carried out with a plan-apochromat 60x or 63x/1.4NA oil-immersion objective. Low magnification images were acquired with a plan-apochromat 20x/0.75NA oil-immersion objective. Pixel size and Z-step were set to fulfil Nyquist criterion. In all cases, lasers and spectral bands were chosen to maximize signal recovery while avoiding signal bleed-through.

Post-imaging analysis and measurements were done using Fiji-ImageJ software (NIH, USA).

### Stimulated Emission Depletion Nanoscopy (STED)

Stimulated Emission Depletion nanoscopy (STED) was performed in a custom-built nanoscope at the Center for Bionanoscience Research (CIBION, CONICET). A detailed description of the setup was provided in a previous publication (Szalai et al., 2021). Briefly, two linearly polarized lasers pulsed at 640 nm (200 ps pulse width, PicoQuant LDH-P-C-640B) and at 594 nm (100 ps pulse width, PicoQuant LDH-D-TA-595) were used for fluorescence excitation, both operating at 40 MHz. In order to obtain circular polarization, both beams passed through broadband (400-800 nm) quarter-wave plates (Thorlabs AQWP05M-600) and broadband (400-800 nm) half-wave plates (Thorlabs AHWP05M-600).

For depletion, a linearly polarized laser pulsed at 775 nm (800 ps pulse width, Onefive Katana HP) operating at 40 MHz was used. The depletion beam passed through a 2π vortex phase plate (Vortex Photonics, V-775-70) in order to obtain a doughnut-shaped beam. Circular polarization was obtained using a broadband (690-1200 nm) quarter-wave plate (Thorlabs AQWP05M-980) and a broadband (690-1200 nm) half-wave plate (Thorlabs AHWP05M-980). The three lasers were combined using suitable dichroic mirrors, and light was focused on the sample with an objective of 1.4 NA (Leica HCX PL APO 100x/1.40-0.70 oil CS).

Fluorescence arising from the sample was split with a long-pass dichroic mirror (FF649-Di01-25x36, Semrock). The reflected light passed through a band-pass filter (FF01-623/24-25, Semrock) and was focused on an avalanche photodiode (APD, SPCM-AQR-13, PerkinElmer Optoelectronics). The transmitted light passed through a band-pass filter (FF01-676/37-25, Semrock) and was focused on a second avalanche photodiode detector (APD, SPCM-AQR-13, PerkinElmer Optoelectronics).

The depletion beam was delayed 0.4 ns with respect to the excitation pulses, in order to maximize the depletion. Time gating of fluorescence photons was also performed, using a custom-made electronic board (MPI for Biophysical Chemistry).

The power of the lasers was adjusted to obtain the best possible resolution in a given set of stained samples, and the imaging conditions were maintained within samples of the same experiment. The optimum values finally obtained were 3 μW and 1.5 μW for 594 nm and 640 nm, respectively. For the depletion beam, a laser pulse energy of 5 nJ was used.

The lateral resolution reached was 50-60 nm in both detection channels. This value was obtained by imaging isolated single molecules and measuring the full width at half maximum (FWHM) of the obtained images.

### Ratiometric FRET Analysis

Fluorescence resonance energy transfer (FRET) biosensor were transfected using lipofectamine 2000 as described above. After 10 to 16 h, cell cultures were fixed with 4% paraformaldehyde - 4% sucrose in PBS for 20 min, washed three times (4 min each) in PBS and mounted in a microscope slide with FluorSave. Cells were visualized with a FV300 or FV1000 confocal Olympus microscopes or with an Olympus IX81 inverted microscope equipped with a total internal reflection fluorescence Cell^TIRF module. For FRET detection, the cyan or teal fluorescent protein (FRET donor) was excited with a continuous laser of 458 nm while simultaneously acquiring donor and acceptor (yellow or venus fluorescent protein) emissions signals. For ratio imaging FRET calculation, donor channel (donor emission) and FRET channel (acceptor emission) images were smoothed with a median filter (1.5 pixel ratio), background subtracted (50.0 pixels rolling back radius) and aligned. FRET map images where generated by dividing the processed FRET channel images over donor channel images. To remove from the analysis out-of-cell pixels, a 0 - 1 intensity values, binary mask was created using the FRET channel images and multiplied by the FRET map images (Quassollo et al., 2015; Rozes Salvador et al., 2016). Finally, pixel values of the FRET maps images were color coded using a custom look up table (LUT). All image processing was coded in a ImageJ macro so all images were processed in the same way.

### Acceptor photo-bleaching FRET analysis

Neurons were transfected with FRET biosensors with lipofectamine 2000 as described above. At the time of analysis, neurons were fixed and mounted in a microscopic slide. Pre-bleaching CFP (donor) and YFP (acceptor) images for Rho biosensors were acquired as described for ratiometric FRET. Acceptor photo-bleaching was done by continuous ilumination of the sample with the 515nm lasers at maximal power until the YFP signal was at least 20% of its original value. A donor and acceptor pair of images (post-bleaching) were acquired with the same configuration as the pre-bleaching images. FRET maps were calculated by applying the following equation: FRETmap = 1 - (pre-bleaching donor image / post-bleaching donor image)

All images were smoothed with a median filter (1.5 pixel ratio), background subtracted (50.0 pixels rolling back radius) and aligned before FRETmap calculation. To remove from the analysis out-of-cell pixels, a 0 - 1 intensity values, binary mask was created using the acceptor images and multiplied by the FRET map images.

### Axon definition in FRET analyses

Early stage 3 neurons were defined as those in which we could clearly identify a single axon determined as a neurite which is at least twice as long as all other neurites but shorter than 60 µm. Late stage 3 neurons were defined as those having a clearly defined single axon of 60 µm or longer and at least twice as long as the remaining neurites.

### Morphometric Analysis

Neurons were immuno-stained for Tau1 and tyrosinated tubulin for morphometric analysis as described above and in previous publications (Wojnacki et al., 2021b; Wojnacki et al., 2021a). The maximal intensity projection of all optical slices of each neuron was used for morphometric analysis. Axons were defined as any process longer than 100 µm and Tau1-positive staining. Unipolar neurons were defined as those with only one axon. Multipolar neurons were defined as those bearing two or more axons. Total axonal length was measured as the cumulative lengths of the longest uninterrupted process and all shorter branches of the axon. Minor process’s length was also measured by the cumulative length of the longest neurite and all branches if present. Fiji/ImageJ and the Simple Neurite Tracer plugin were used to measure all neurites lengths (Schindelin et al., 2012).

### Live-cell imaging and EB3 comet analysis

Fluorescent images of living neurons were acquired as previously described (Bisbal et al., 2016). Briefly, transfected neurons expressing EB3-GFP were captured with a charge-coupled device camera (Andor iXon3; Oxford instruments) using an 60x/NA 1.4 oil immersion objective in an Olympus IX81 inverted microscope equipped with a Disk Spinning Unit (DSU) with epifluorescence illumination (150 W Xenon Lamp). Neurons were imaged in neurobasal medium supplemented with 30 mM HEPES buffer (pH 7.2) and maintained at 37 °C. Time-lapse sequences were acquired at a continuous rate of 1 frame per second during 5-10 min inside a stage top incubator (INU series, TOKAI HIT). Images were processed offline using a plusTipTracker from µ-Track software Version 2.0 (https://www.utsouthwestern.edu/labs/danuser/software) (Applegate et al., 2011; Jaqaman et al., 2008). Movies were color coded using a temporal-color code look up table (LUT) to generate an XY 2D image.

### Cortical Scratch assay

Cortical scratch assays were performed as described previously (Kaplan et al., 2017). Briefly, on DIV 9, two parallel scratches (3 mm apart) were made across the coverslip with a plastic P10 pipet tip. Treatments, transfections and fixation for immunofluorescence were performed according to the procedures described in the corresponding sections and at the time-points indicated in the text. Images of βIII-tubulin stained axons were acquired in the center of each scratch and thresholded in ImageJ. The outline of each scratch was traced and, at 8 h, the extent of axon growth was quantified as the total area of βIII-tubulin in the scratch, while at 24 h the extent of axon elongation was measured as the percent area of βIII-tubulin stained neurites in the central 70% of the scratch. Image analyses were performed with ImageJ. All assays were performed with 6-12 replicates per condition from 3 independent experiments.

### Evaluation of Growth cone fluorescent intensity

For the images where the fluorescence intensity was quantified, once the microscope was configured on the first acquired image, the same configuration was maintained for the acquisition of all subsequent images. The maximal intensity projection of all optical slices of each neuron was used for fluorescence intensity analysis Adjusting the intensity level, we excluded from the analysis all pixels with values less than 200 and cells with saturated pixels. Subsequently, the regions of interest (axonal growth cones or minor processes) were selected and the average fluorescence intensity value per region was quantified.

### Evaluation of Growth cone microtubules

For quantification of the number and length of microtubules in growth cones STED images were used. All the images between experiments were acquired with same microscope configuration. Growth cones areas were defined making a region of interest (ROI) in the phalloidin channel. Individual microtubules were quantified within the ROI and the length measured from the base of the growth cone to the tip of the microtubule.

### Statistical analyses

Statistical analysis was performed with the R software and Infostat software. When the experimental design required a single pairwise comparison, Student *t*-tests were applied. If the experimental design required, the comparison of multiple conditions a one-way ANOVA test was used followed by a Turkey’s HSD test. The exact value of n, what it represents and center and dispersion measures are described in the figures and figure legends. In experiments where we measured neurite length or immuno-fluorescence intensity, each cell was considered as an independent statistical observation. Statistical observations come from at least three independent experiments. Significance was defined as a p value lower than 0.05 unless otherwise stated. The lower and upper hinges of all boxplots correspond to the first and third quartiles respectively (the 25th and 75th percentiles). The upper whisker extends from the hinge to the largest value no further than 1.5 * IQR from the hinge (where IQR is the inter-quartile range, or distance between the first and third quartiles). The lower whisker extends from the hinge to the smallest value at most 1.5 * IQR of the hinge. Data beyond the end of the whiskers are called ‘‘outlying’’ points and were plotted individually.

## Author contributions

Conceptualization, A.C., M.B. and J.W.; Formal analysis, J.W, G.Q., M.D.B, A.M.S.; Funding acquisition, A.C. and M.B.; Investigation, J.W, G.Q., M.D.B, N.U., G.F.M., A.M.S. and M.B.; Methodology, J.W, G.Q., M.D.B, N.U., F.D.S., A.C., A.M.S. and M.B.; Project administration, A.C. and M.B.; Resources, O.P., G.G.G., F.B., F.D.S., A.C. and M.B.; Supervision, A.C. and M.B.; Visualization, J.W, G.Q., M.D.B, N.U., A.M.S., A.C. and M.B.; Writing—original draft, A.C.; Writing—review and editing, J.W., O.P., G.G.G., F.B., F.D.S., A.C. and M.B. All authors have read and agreed to the published version of the manuscript.

## Conflict of interest

The authors declare no competing financial interests.

## Supporting information

Wojnacki et al. Supplementary Figures 2023

## Acknowledgments

This research was funded by grants from Agencia Nacional de Promoción Científica y Tecnológica (Argentina), PICT 2015-1436 (A.C.) and by the International Brain Research Organization, IBRO Early Career Awards 2020 (M.B.). N.U., F.D.S., A.C. and M.B. are staff scientists from the National Council on Scientific and Technical Research (CONICET). The authors greatly acknowledge the Centro de Micro y Nanoscopía de Córdoba (CEMINCO), for technical and imaging assistance. nt financial or non-financial interests to disclose.

## Data availability

The datasets generated and/or analyzed during this study are available from the corresponding author on request.

## Notes

### Competing Interest Statement

The authors have declared no competing interest.

### Summary of Updates

Figure 4 revised. Figure 6 revised with new experiments

