## Supplementary material for "Dual spatio-temporal regulation of axon growth and microtubule dynamics by RhoA signaling pathways": Wojnacki et al. Supplementary Figures 2023

***Abbreviated title*: *Dual regulation of axon growth by RhoA signaling.***

José Wojnacki[1†], Gonzalo Quassollo[1†], Martín D. Bordenave[3], Nicolás Unsain[1, 2], Gaby F. Martínez[1], Alan M. Szalai[3], Olivier Pertz[4], Gregg G. Gundersen[5], Francesca Bartolini[5], Fernando D. Stefani[3, 6], Alfredo Cáceres[7*], Mariano Bisbal[1, 2*]

**Affiliations:**

[1]Instituto de Investigación Médica Mercedes y Martín Ferreyra (INIMEC), Consejo Nacional de Investigaciones Científicas y Técnicas (CONICET), Universidad Nacional de Córdoba, Córdoba, 5016, Argentina.
[2]Instituto Universitario Ciencias Biomédicas de Córdoba (IUCBC), Córdoba, 5016, Argentina.
[3]Centro de Investigaciones en Bionanociencias (CIBION), Consejo Nacional de Investigaciones Científicas y Técnicas (CONICET), Godoy Cruz 2390, Ciudad Autónoma de Buenos Aires, C1425FQD, Argentina.
[4]Institute of Cell Biology, University of Bern, Baltzerstrasse 4, Bern, 3012, Switzerland.
[5]Department of Pathology and Cell Biology, Vagelos College of Physicians and Surgeons, Columbia University, New York, NY 10032, USA.
[6]Departamento de Física, Facultad de Ciencias Exactas y Naturales, Universidad de Buenos Aires, Güiraldes 2620, Ciudad Autónoma de Buenos Aires, C1428EHA, Argentina.
[7]Centro Investigación Medicina Traslacional Severo R Amuchástegui (CIMETSA), Instituto Universitario Ciencias Biomédicas Córdoba (IUCBC), Av. Naciones Unidas 440, Córdoba, 5016, Argentina.

†These authors contributed equally to this work.

*To whom correspondence should be addressed:

Alfredo Cáceres

ORCID 0000-0002-4163-9068

Mariano Bisbal

ORCID 0000-0002-3870-6151

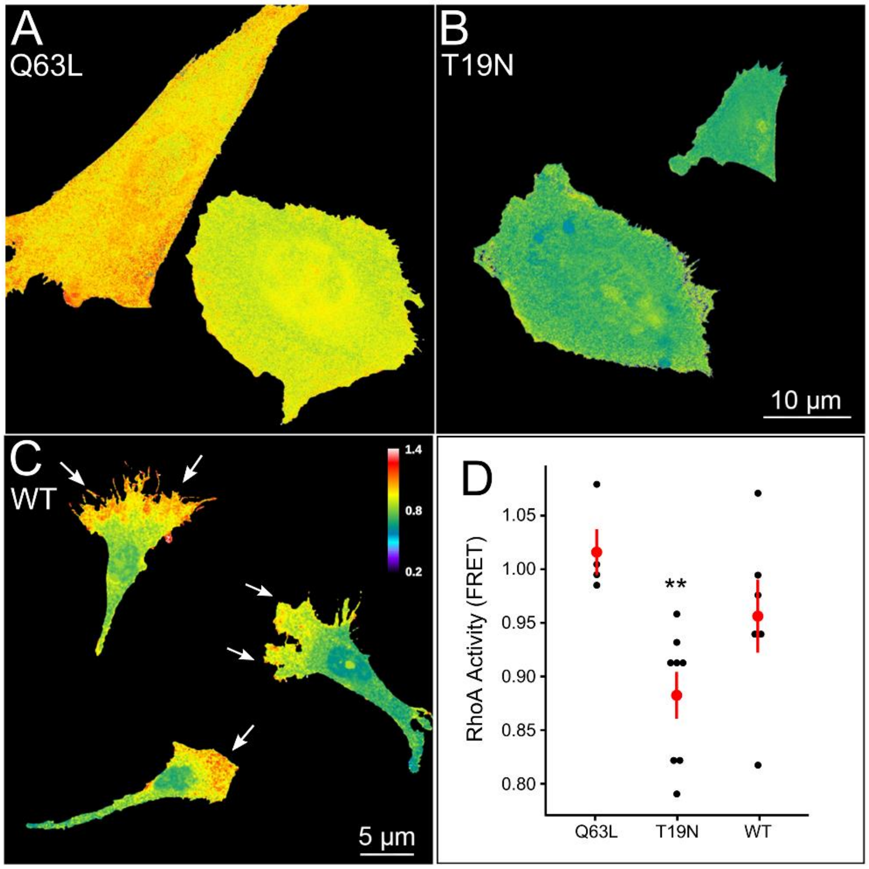


**Supplementary Figure 1. RhoA activity radiometric FRET assay characterization**

(*A*, *C*) Representative FRET maps images of CHO cells expressing the unimolecular FRET biosensors for the RhoA mutants: RhoA Q63L (constitutive active, *A*); RhoA T19N (dominant negative, *B*) and RhoA wildtype (*C*). Radiometric method was used to measure the FRET signal and FRET maps are color-coded according to activation intensity. Color-code according to the scale bar. Note that the highest RhoA FRET signal at the leading edge in wildtype biosensor (arrows). (*D*) Graph showing the quantification of the mean activity of the different RhoA mutants calculated by radiometric FRET. Each black dot represents the average FRET value of a single CHO cell. Each large red dot represents the mean ± S.E.M.; **<0.01; one-way ANOVA and Tukey's *post hoc* test. Cells were obtained from two independent cell cultres.

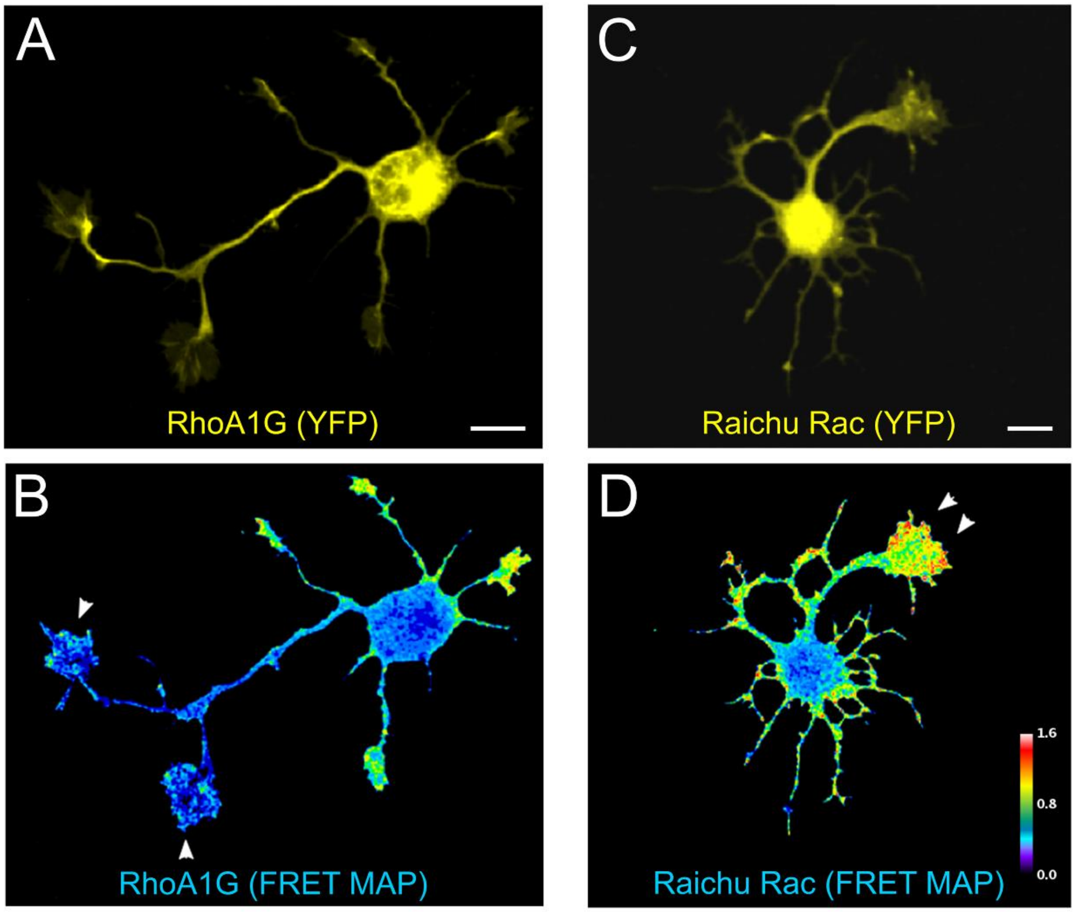


**Supplementary Figure 2. Rho GTPases activity during axon differentiation**

($A$, $B$) Representative image and the corresponding FRET map showing the distribution ($A$) and activity ($B$) of the FRET biosensor RhoA1G in a 1 DIV cultured hippocampal neuron shortly after transitioning from stage 2 to stage 3. Scale bar: 10 µm ($C$, $D$) Representative image and the corresponding FRET map showing the distribution ($C$) and activity ($D$) of FRET biosensor Raichu Rac1 in a 1 DIV cultured hippocampal neuron transitioning from stage 2 to stage 3. Scale bar: 10 µm. Radiometric method was used to measure the FRET signal and the FRET maps are color-coded according to activation intensity. Color-code according to the scale bar.

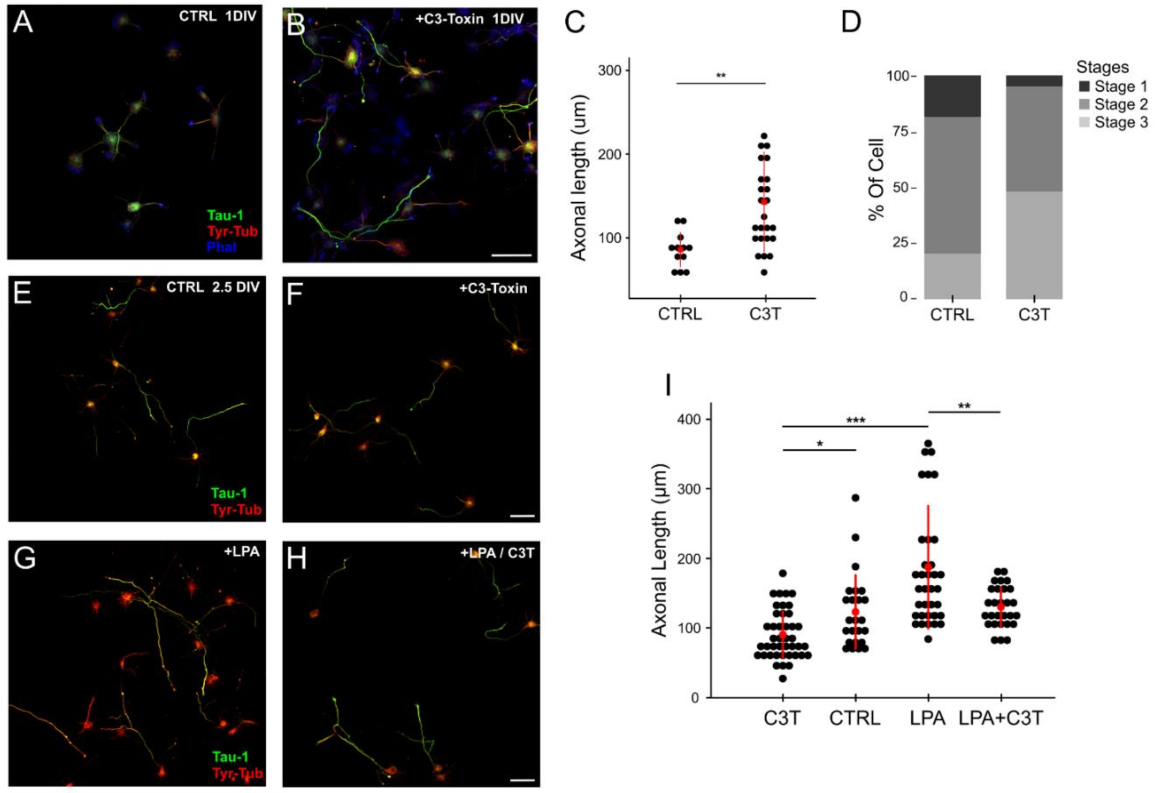


**Supplementary Figure 3. RhoA have opposite effects in axonal growth during neuronal polarization**

(*A*, *B*) Representative confocal images of cultured hippocampal neurons (1 DIV) treated with vehicle (DMSO, *A*) or with the RhoA inhibitor C3-toxin (0.5 µg/ml, *B*) for 24 h, fixed and stained with the mAb Tau-1 (green), Ab Tyrosinated tubulin (red) and Phalloidin-Alexa 633 (blue). Scale bar: 50 µm. (*C*) Graph showing quantification of the axonal length of figures A and B. Each black dot represents the axonal length of a single neuron. Red dots represent the mean total axonal length ± S.E.M.; *<0.05; one-way ANOVA and Tukey's *post hoc* test. For all experiment, 14 to 24 neurons were quantified pooled from at least three independent cultures. (*D*) Graph showing the percentage of stages 1, 2 and 3 in (*A*, *B*). The percentages were calculated from 3 independent cultures. (*E*-*H*) Representative confocal images of cultured hippocampal neurons (2.5 DIV) treated with vehicle (DMSO, *E*), C3T (0.5 µg/ml, *B*), LPA (10 µM, *F*) or LPA + C3T (10 µM / 0.5 µg/ml, *H*) for 24 h, fixed and stained with the mAb Tau-1 (green) and Ab Tyrosinated tubulin (red). Scale bar: 50 µm. (*I*) Graph showing quantification of the axonal length of figures *E*-*H*. Each black dot represents the axonal length of a single neuron. Red dots represent the mean total axonal length ± S.E.M.; *<0.05; one-way ANOVA and Tukey's *post hoc* test. For all experiment, 25 to 43 neurons were quantified pooled from at least three independent cultures.

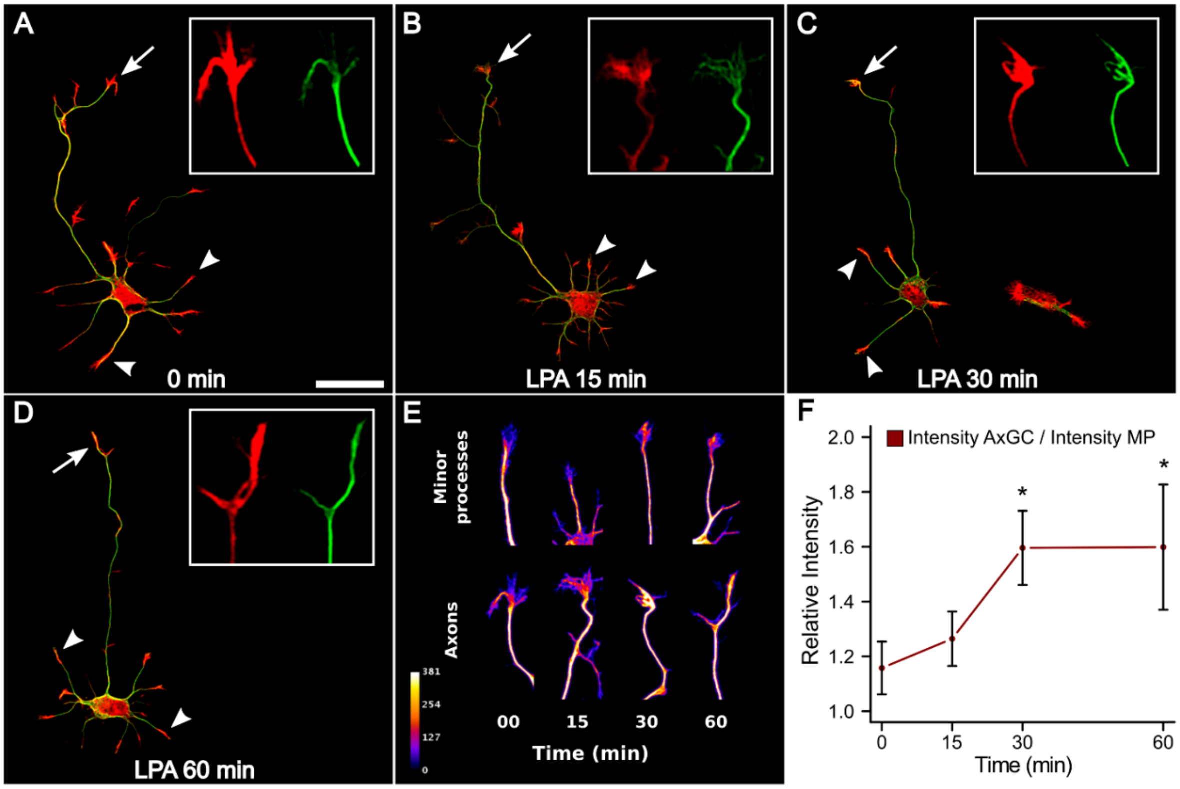


**Supplementary Figure 4. LPA induced an increase in acetylated tubulin fluorescence in axonal growth cones during axonal elongation**

(*A-D*) Representative confocal images of cultured hippocampal neurons (3 DIV) treated with LPA (2.5 μM) for 0, 15, 30 and 60 minutes respectively. Neurons were stained with mAb acetylated α-tubulin (green) and phalloidin-rhodamine (red). Scale bar: 20 μm. Insets show single channel high magnification views of axonal growth cones. The intensities of both channels are on the same scale in all images. (*E*) Acetylated α-tubulin channel of the axonal growth cones (arrows in *A-D*) and minor processes (arrowheads in *A-D*) shown in *A-D* assigned to a pseudo-color that reflect differences in fluorescence intensity (Fire LUT, color code bar). (*F*) Graphs showing the quantification of the acetylated α-tubulin fluorescence intensity at the different times point of treatment with LPA. The intensity of fluorescence in the axonal growth cones is expressed in relation to the intensity of the minor processes (red). Dots represent the mean acetylated α-tubulin intensity ± S.E.M.; *<0.05; one-way ANOVA and Tukey's *post hoc* test. Asterisks mark significant differences with respect to time 0 minutes. For all experiment, 12 to 18 neurons were quantified pooled from at least three independent cultures.

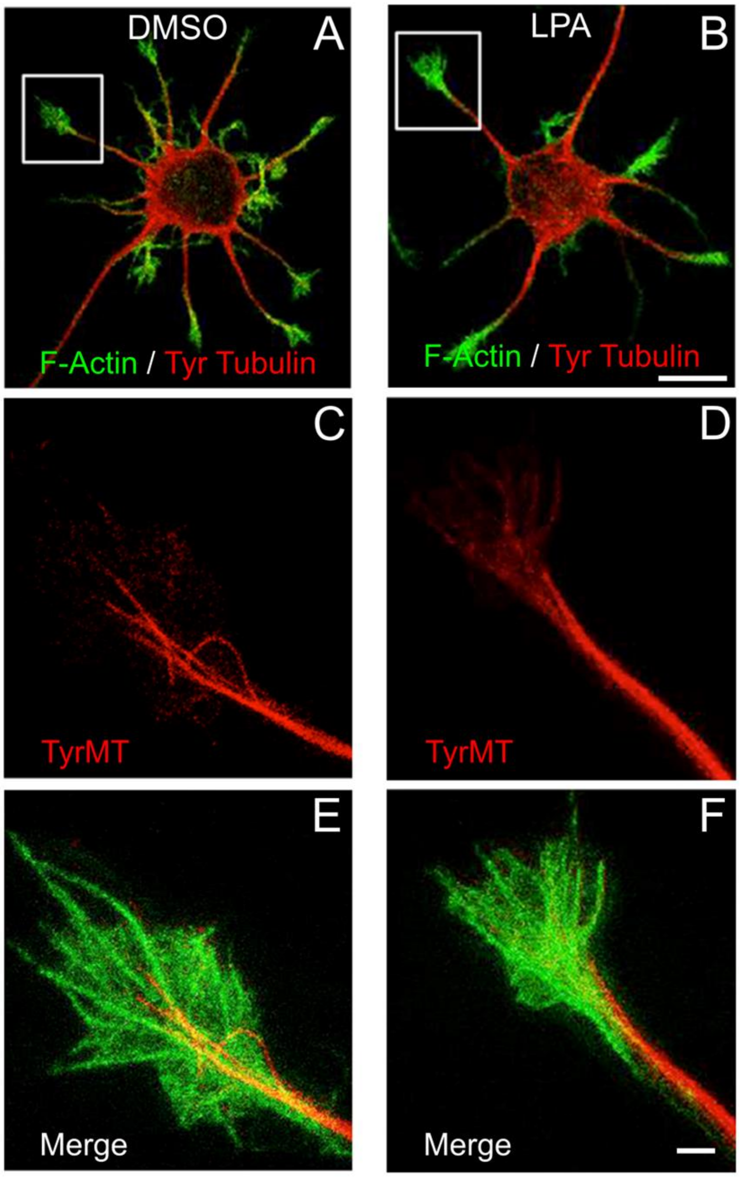


**Supplementary Figure 5. LPA does not promote microtubules abundance in minor process growth cones.**

(*A*, *B*) STED microscopy images showing representative 3DIV cultured hippocampal neurons treated with vehicle (DMSO, *A*) or 10 µM LPA (*B*) stained with mAb tyrosinated α-tubulin (red) and phalloidin-atto594 (green). Scale bar: 10 µm. (*C*-*F*) High magnification views of the inserts shown in *A* and *B*, showing a representative minor process growth cone. Scale bar: 2 µm.
